# Adaptive learning under strategic and structural uncertainty: the case of auction games

**DOI:** 10.1101/2020.08.22.262469

**Authors:** Mario Martinez-Saito, Alexis Belianin, Anna Shestakova, Boris Gutkin, Vasily Klucharev

## Abstract

In games of incomplete information individual players make decisions facing a combination of structural uncertainty about the underlying parameters of the environment, and strategic uncertainty about the actions undertaken by their partners. How well are human actors able to cope with these uncertainties, and what models best describe their learning in such environments? We use a double auction task with different competitive and informational environments to characterize learning abilities of the single human participants (buyers) in a range of adaptive learning models covering reinforcement learning, directional learning and belief learning. Results show that real behaviour is best described using simple models of directional learning type with minimal knowledge assumptions about information efficiency of prices. This behavior is consistent with bounded rationality and risk aversion: human subjects try to maximize their chance for transaction, and do so using the simplest learning rule.

## 1. Introduction

How do people integrate past information to assign value to goods which are subject to changing strategic player interactions? Learning about how people play interactive play games has been an important topic in economics for quite some time (Roth and Erev, 1995; Fudenberg and Levine, 1998; 2016). nNumerous learning models, such as fictitious play or reinforcement learning have been proposed and successfully applied in a variety of contexts (Erev and Roth, 1998; Binmore and Samuelson, 1998; Weibull, 1995; Forster and Vohra, 1997; Hart and MasColell, 2000, Abbink and Cox, 2005). Yet in other cases, these same models have not been very accurate descriptively, especially if players face uncertainty about the expected outcome or underlying parameters of the game, or incentives, preferences, beliefs or reasoning procedure used by their opponents. All these factors may be attributable to two potential sources (Harsanyi and Selten, 1988; Brandenburger, 1996; Andersson et al., 2012): *structural uncertainty* about the characteristics of the environment, and *strategic uncertainty* about preferences, beliefs and behavioral strategies of the other players. If either of these characteristics are not commonly known, human players have to form their own representation of the problem, and react according to it.

In our paper we study human behaviour in face of structural and strategic uncertainty, and a combination of these, which is typical in many real market interactions. As a prototype game, we take the double auction games with exogenously varying number of buyers and sellers. These games are very well explored in the literature on bargaining and experimental markets, and are shown to result in remarkably robust behaviour, which is well-aligned with theoretical predictions. In these canonical market environments, we vary information about the true market type to study the impact of structural uncertainty, and information about other players’ decisions to study strategic one. In a nutshell, our results show that human subjects respond to these uncertainties by lowering their bids to increase the probability of a profitable deal, at a cost of decreased expected profit. Finally, we fit the observed strategies to the various learning models, and explore comparative predictive power of learning strategies in these market environments. Estimates of these models favour the simplest possible learning rule – Directional learning. This fact may be viewed as evidence of ‘economic’ response to the complexity of decision task whenever the cost of using more involved learning strategies fails to overweight their expected benefits.

Our variation of the market environments induces natural variation in decisions. For instance, if a buyer repeatedly competes with another buyer over a good offered by a single seller, she will soon realize that in order to avoid the loss of a profitable deal, she need to increase her offer. This situation contrasts with the opposite case, in which of two or more sellers compete the demand of a single monopsonist. In our paper we consider three market games: buyer’s competition (BC) of two symmetric buyers facing a single seller; sellers’ competition (SC), with two symmetric sellers facing one buyer, and neutral competition (NC), which is simple bilateral bargaining of a single buyer with a single seller. Experimental evidence (Huck et al., 1999, 2000; Duwfenberg and Gneezy, 2000; Offerman et al., 2002), including the one reported below, suggests that under full information, human players are relatively quick learners of optimal behavior in each of these market types, which eventually converges to Bertrand Nash equilibrium. A third market type is bilateral bargaining of a single buyer against a single seller, or neutral competition (NC). Unlike the previous case, this situation is a coordination game, akin to Nash (1953) demand game, and yields multiple equilibria laying on the main diagonal. Existing literature (see Ochs, 1995 or Crawford, 1990) suggests that human subjects in this context will coordinate on focal solutions, in case of uncertainty giving greater share to the player who is best informed (Andersson et al., 2018). Comparison of the behavior in games with single and multiple equilibria allows us to control for the effect of strategic complexity on the speed and efficiency of learning.

To obtain a clean picture, throughout the paper we concentrate on the behavior of a single player – the buyer. To ensure constancy of experimental conditions, human buyers were playing against pre-recorded opponents (buyer and seller(s), depending on the actual market type). To capture strategic and structural uncertainty, we use three exogenously varying information conditions. In our *benchmark treatment* (T1) the information about the market type is correct and truly communicated to the buyer, who observes only own outcome, but receives no feedback about what other players did in the previous periods. This situation corresponds to no structural uncertainty with strategic uncertainty in place. In another *“scrambled” treatment* (T2), we exogenously distort this information about market type by scrambling the market labels: instead of the true market type, subjects were presented with randomly assigned market type labels (e.g., SC when the actual market type is BC). Here the buyer still observes the outcome, but not necessarily the true market type, thus creating structural uncertainty. Under these conditions, naive buyers who react only to a signal about the market type may continue playing the strategy corresponding to each pre-announced type, while rational players should respond to factual outcomes, and adjust their behavior to the extent of their learning abilities. Finally, in the *full information treatment* (T3), all market types are correctly announced, and all opponents’ bids are revealed to the subject (buyer). This treatment entails the situation of lowest strategic uncertainty, limited to occasional irregularities in the behavior of pre-recorded partners. This environmemt is expected to result in most efficient learning.

Each subject participated in all three treatments, presented to participants in random order. All treatments consist of 24 replications, which number is reasonably short to prevent subject from being exhausted, but sufficiently large to learn in each market type. Comparing buyers’ behavior under the various information conditions in each of the three market types, we can disentangle the effect of strategic uncertainty (baseline vs full information treatment) from that of structural uncertainty (full information vs. scrambled treatment).

Our specification of strategic and structural uncertainty has several real-life (ecological) motivations. With some notable exceptions (e.g. financial markets), full information about the behavior of one’s competitors is often not available and/or prohibitively costly. Markets with scrambled information are also more typical in reality than it might appear. Online auctions such as eBay, feature several buyers bargaining for a single unit, without knowing the number of other contestants for the same good. Since all competition is online, the illegal, but still present practice of *shill bidding* from artificial accounts serves to artificially intensify competition and push the price up. In the government procurement auctions, the auctioneer who organizes a tender bid has to set the highest reservation price for the contact, without knowing ex ante how many competitors will come. Hence, a single potential bidder has strategic incentives to pretend he will be the sole competitor. In fact, this may well not be true, but this impression could persuade the auctioneer to increase the reservation price in order not to lose the supplier. In such cases, scrambled information about market structure may be part of the bargaining strategy of one of the sides of the transaction. Our experimental design mimic such situations, allowing us to compare the efficiency of coping strategies of real players in such adverse information conditions.

These strategies are crucially related to learning in various strategic environments (Erev and Haruvy, 2015). Standard economic reasoning suggests that people will respond to incentives, and converge to equilibria in the respective games, although at a different rate, depending on the accuracy of information available in our various treatments. The BC and SC market environments the unique predictions are in favour of the parties with more bargaining power, i.e prices equal to the upper bound of the range (buyers’ valuation) in the SC and to the lower bound (seller’s costs) in the BC. In the non-cooperative NC game, there is a continuum of possible equilibria, but invoking the symmetry axiom, we would predict convergence to the Nash bargaining solution and equal division of the joint surplus. See Appendix C for the details).These predictions receive empirical confirmation in our data. In particular, subjects learn in the predicted directions, and do so most efficiently in the full-information treatment. In the non-scrambled treatment, buyers’ strategies also follow the predicted patterns, converging to the Bertrand Nash equilibrium in BC and SC, and the Nash bargaining solution in the NC market type, respectively. In T2 (scrambled treatment), we find that the buyers’ bids are systematically biased upwards in all market types. This finding can be explained by risk aversion of the buyer who is afraid of not making a deal (see Appendix C for the theoretical argument).

Finally, we compare the explanatory power of several alternative learning models in all three markets and all informational treatments, including directional learning, belief learning and reinforcement learning. After fitting the model parameters by maximum likelihood, we find that the best is the simplest one – *directional learning model*. Altogether, these findings are in line with bounded rationality theory: players in an uncertain strategic situation strive to maximize their chance for transaction, and do so using the simplest learning rule.

The rest of the paper is organized as follows. Section 2 puts our experimental design into perspective with respect to the relevant literature. Section 3 explains the setup of our experimental game. Section 4 presents the six learning rules that have been applied to our experimental data, and section 5 presents the results. Section 6 discusses the relevance and importance of our findings. Section 7 concludes.

## 2 Related literature

Studies of learning dynamics in games are not new to literature in economics and computer science, including the settings of auction games. The mainstream literature focuses on learning how to play Nash equilibrium (or a specified subset of these, if there is more than one). The primary justification for Nash equilibrium is evolutionary (Nash, 1951; Weibull, 1995). Indeed, much of the existing experimental evidence suggests that strategies of real subjects may start quite away from Nash equilibrium, but gradually converge to it (Cheung and Friedman, 1997; Binmore and Samuelson, 1997; Fudenberg and Levine, 1998). In the simplest case of normal-form game, subjects observe only the payoffs collected in the previous stage, and adjust own strategies given the experience with the others. This yields learning dynamics of best reply type, such as fictitious play (Brown, 1951) and stochastic fictitious play (Fudenberg and Kreps, 1993). These models assume that real subjects believe their opponents play some fixed but unknown mixed strategy, augmented in stochastic version with random payoff shocks, and play best reply against an empirical sequence of opponents’ plays, where probabilities of each expected play is updated in accordance with the Bayes rule. This scheme is arguably very rigid: in particular, it may result in strategy cycles, which are not always easy to accommodate (Aoyiagi, 1996), and is unable to incorporate some forms of strategic experimentation (e.g. playing best replies to strategies with zero posterior probability - Fudenberg and Levine, 2009). These facts have called for a broader perspective of learning. In particular, players may be Bayesians when re-weighting the upcoming evidence about types of the others, but their priors about these types need not necessarily correspond to the ones, which lead to Nash equilibrium. However, any particular prior coupled with Bayesian updating and based on calibrated forecasts leads to correlated equlibria (Foster and Vohra, 1997). This finding suggests to look at bounded rational leading strategies based on simple clues: Hart and Mas-Colell (2000) argue that learning inertia and regret (of not playing the action which proved to be good) almost surely converges to a correlated equilibrium, but not to a Nash equilibrium. Hart and Mas-Collel (2003) show that this last property holds for all uncoupled dynamics, which are generic to boundedly rational rules in the sense that learning players may observe the strategies, but not payoffs to the others, as in our benchmark treatment T1. This goes in contrast with the classical learning rules, such as replicator dynamics (Taylor and Jonker, 1979), stochastic fictitious play (Fudenberg and Levine, 2009) and quantal response (McKelvey and Palfrey, 1995). These models, under various conditions, all asymptotically converge to Nash, although the speed of convergence remains a separate, and largely underexplored issue (Fudenberg and Levine, 2016). te, and largely underexplored issue (Fudenberg and Levine, 2016).

Decisions under strategic uncertainty has been studied both experimentally (Abbink and Brands, 2008; Feltowich and Swierzbinski, 2011; Andersson et al., 2018) and theoretically (Andersson et al., 2014, Yamashita, 2015). Some works, including Abbink and Brandts (2005) studied the behavior under structural uncertainty, although most of the literature focused on pure effects of the varying market structures (Plott, 1995, Camerer, 2003 and Huck et al. 2004 survey the relevant literature).

A separate branch of literature explores convergence in the specific settings of bargaining. Rapoport, Daniel and Seale (1998a; 1998b; 2000, 2001), in a series of experiments explored the predictive power of the standard double auction game with incomplete information (Chatterjee and Samuelson, 1983; Radner and Schotter, 1989; Myerson and Satterthwaite, 1989; Linhart et al., 1992), and found that the theoretical prediction of symmetric linear Bayesian Nash equilibrium works reasonably well for under various valuation, information and treatment conditions, especially for the sellers, and less so for the buyers. However, these authors did not explore bargaining behavior under changing market structures. Furthermore, Croson et al. (2003) have shown that subject strategies in incomplete information bargaining games with cheap talk can be quite complicated, and involve strategic misleading. Extensive experiments in auction games (Kagel, 1995), while confirming consistency of the lab evidence with the theory, reveals that learning is sometimes slow, especially in low-information, sealed bid treatments (Kagel et al., 1987; Ariely et al., 2005). Extensive experimental studies of various market structures (see Plott, 1989 for a classical survey), confirm the ultimate effect of market power on outcomes, but do not offer a concise story of convergence to market equilibria. For example, Kutschinski et al. (2003) in a field study based on electronic auctions demonstrate that joint learning of several adaptive agents may lead to fuzzy dynamics that is both unforeseen and increasingly volatile.

This literature suggests that the logic of Nash equilibrium, while remaining a strong attractor in some contexts, may fail descriptively for a range of behavior which may nevertheless be quite rationalizable on alternative grounds. One way to relax the assumptions of Nash is to forego belief modeling (which are typically unobservable anyway), and concentrate on immediate impulses from experienced utilities (payoffs). This approach of reinforcement learning had emerged in natural sciences, and has been introduced in economics by Roth and Erev (1995; 1999), and further developed in the experience weighted attraction (EWA) model (Camerer and Ho, 2000). It is worth notice, however, that this approach is by no means exclusive nor universal to all contexts. In fact, most models fall short of explaining the underlying generating process by which human players learn in economic decision tasks (Salmon, 2001). Selten, Chmura and Goerg (2011) compare descriptive performance of several models of learning against an empirical dataset based on more than 800 subjects playing for 200 rounds. They find that simple heuristic strategies, such as impulse balance equilibrium (a concept coined by Selten in 1998, based on aspiration levels of the players) as well as payoff and action-sampling equilibrium (Osborne and Rubinstein 1998) outperform equilibrium models, such as quantal response equilibrium (McKelvey and Palfrey, 1998) and Nash equilibrium. Simple rules also outperform more fundamental learning rules, such as linear and logarithmic tracing procedures (Harsanyi, 1974; Harsanyi and Selten 1988). These latter are widely used to find the set of Nash equilibria in an arbitrary finite game in strategic form (see von Stengel, 2002 for a review). despite logical correctness and formal attractiveness, actual behavior no more corresponds to these rules than does their behavior converge to Nash equilibria.

To sum up, the literature suggests that even in simple games with perfect ex ante information but uncertain environment and/or preferences of the other players, Nash equilibrium and its direct generalizations, such as quantal response or fictitious play, are not among the best predictors to actual behavior. By contrast, simpler, boundedly rational learning rules better perform descriptively, and correspond better to the set of rationalizable decisions in ecologically relevant conditions. These include unobservable priors about opponents’ types, correlated equilibria, and covering strategic manipulation of information about the institutional environment upon which one has to act. Analysis of behavior in such settings calls for a controlled experimental study, to which we proceed now.

## 3. Materials and Methods

### 3.1 Experimental paradigm

Our market (double auction) experimental game was framed as the problem of a single trader (restaurant owner) who aims at buying a fixed-size commodity (fish) to be sold to his or her customers at a fixed price, and follows the design of a previous study (Martinez-Saito et al., 2019). The trader in each period comes to the market, where he faces one of the three possible market types: SC (two sellers, each having and willing to sell him one unit of good), NC (single seller with one unit of good), or BC (single seller but another buyer, competing with the trader for the same unit of good). Only the trader is actual participant of the experiment; other agents are dummy players whose decisions are pre-recorded and displayed automatically. In a computerized game, the trader plays the role of a buyer facing market types presented sequentially in random order. At the beginning of each trading period, each buyer has a fixed endowment to buy fish (at the cheapest price possible).

The following parameters have been used for the experimental game. Buyer valuation always equals 10 monetary units (MU), and seller’s valuation (cost) is always normalized to 0. The active buyer (player) *b1* decides his bid; opponents’ decisions of all other players (single seller *s1* in NC, two sellers *s1* and *s2* in SC and seller and other buyer *b2* in BC) were drawn from the previous experimental data (Muller, 2013) by random-opponent matching (see Figure 2 for the presentation of this result).

Each trial consisted of three stages. In the first stage, which lasted for 5 seconds, the player is informed about the market type (s)he will face in the current trial (see below). In the second, the player had up to 15 seconds to make their bids *b* by placing a bar at a proper point on a scale from 0 to 10. At the final stage, player’s decision is matched with that of the other agents, and outcome of the trial is displayed for 6 seconds on the computer screen. The feedback screen contained information about the transaction outcome (accepted or rejected), the payoff earned by the subject, as well as the bid of the second buyer in the BC condition. Figure 1 shows the sample sequence of screen stages, programmed using Presentation software (version 18.0, Neurobehavioral Systems, USA, www.neurobs.com).

**Figure 1.**
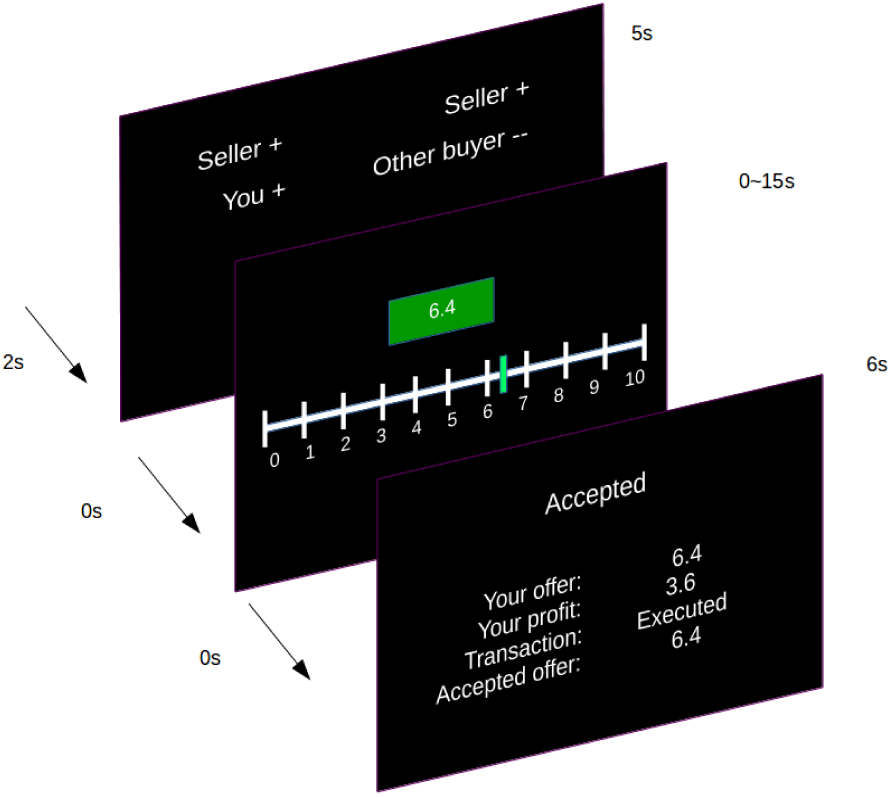
Task flow. Each trial consisted of three stages: market type announcement, bid selection, and game outcome feedback. During the market announcement stage, the subject is informed of the market type of the current trial. Next a scale from 0 to 10 is displayed, and the subject has to choose her bid by sliding a vertical bar. Finally, at the transaction outcome stage, trade outcome and payoffs to the player is announced, along with an information as to whether the bid was accepted or rejected.

Instructions for the game given to the subjects (English version) are contained in Appendix B.

### 3.2 Treatments

Our study consisted of 3 treatments (see Table 2 below for a summary). The baseline *benchmark treatment* (T1) has been conducted exactly as described above, with subjects knowing only their own performance. To explore the role of information, we add two more treatments. Additionally, we conducted two *control treatments* (T2 and T3) defined a follows. In T2, the market type announcement at the beginning of each trial was random instead of the actual one. That is, we uniformly randomized the three labels (SC, NC, BC markets types), so that the signal was scrambled, and subjects played in the announced market type only 1/3 of trials. To ensure parallelism of the treatments, no further information has been ever conveyed to the subjects in all information treatments.

**Table 2.** Number and shares (in parentheses) of successful transaction by experimental treatment and market type. In all treatments and market types, N=432, total number of observations per market type is N=1296.

| Treatments \ Market types (per treatment) | SC (over 432) | NC (over 432) | BC (over 432) | Total (over 1296) |
| --- | --- | --- | --- | --- |
| T1 | 383 (88.7%) | 323 (74.8%) | 224 (51.9%) | 930 (71.8%) |
| T2 | 264 (61.1%) | 277 (64.1%) | 313 (72.5%) | 854 (65.9%) |
| T3 | 387 (89.6%) | 324 (75.0%) | 219 (50.7%) | 930 (71.8%) |

Finally, as a control treatment needed to assess the impact of full feedback on subjects’ behavior, we use T3, where the information about market type was not only correct (as in T1), but also full, i.e. supplemented by all buyer’s bids in every trial.

As robustness checks to these treatment effects, we endow our design with several new features. First, to focus attention on buyers’ bidding strategy, and mitigate potential perceptive errors on the part of the subjects, in the BC markets we set the price of successful transaction at buyers’ bid instead of the usual Nash bargaining price equal to midway between buyer and seller’s proposals. Second, as additional robustness check for reaction to strategic uncertainty, we augment the design with a parameter that is orthogonal to market structures. Namely, in the SC market, when both sellers can make a deal, the one who will actually transact has been determined not by the best (lowest) bid but at random. In the T3 treatment, buyers know both winning and losing bids, hence can form a clearer idea about the strategies of the sellers, which is impossible to do in the non-perfect information treatments (T1 and T2).

Experimental sessions were conducted in March-June 2015. Participants were students, mostly from psychology department in Moscow, recruited via internal recruitment system. Altogether, 54 subjects played 24 blocks of each treatment (T1, T2 and T3), alternating among market types in the same randomly predetermined order throughout the session. Although the alternation order was fixed within each experimental session, it was randomly varied across subjects.

All experiments were conducted separately for each experimental subject who was alone in the room with the computer. After arriving to the classroom and signing the consent form, each participant read the instruction (see Appendix D), and had an opportunity to ask any questions. Subjects were explicitly informed that in all simulated markets, their opponents were prerecorded humans playing in concomitant trials (i. e., the actions of their opponents were matched according to the trial order of each market type), and in the same format (that is, repeated random matching) against other human opponents, and hence in no way different from real humans except that this prevented using strategies to influence opponents’ play (Camerer et al., 2002).

Prerecorded data (Figure 2) were obtained from a pilot study (Muller, 2013). The design of the pilot study task was identical, to the main experiment with the following exceptions: 16 subjects played in anonymous groups; on desktop computers with conventional keyboards were used, and subjects played against each other, simultaneously, within the same room, but uninformed of who their opponents were on a given trial. Subject roles were randomly assigned to buyer *or* seller throughout the duration of the experiment. Both seller and buyer had to set their respective ask prices and bids beforehand. The total number of trials amounted to 11520 (3 sessions, one per market type, each consisting of 40 periods with 6 rounds per period). Bids/ask prices statistics in BC were 7.67+-1.12 for buyers and 3.84+-2.63 for sellers: in NC, 3.27+-2.19 for sellers; and in SC, 2.49+-1.70 for sellers (+-stands for standard error of the mean).

**Figure 2.**
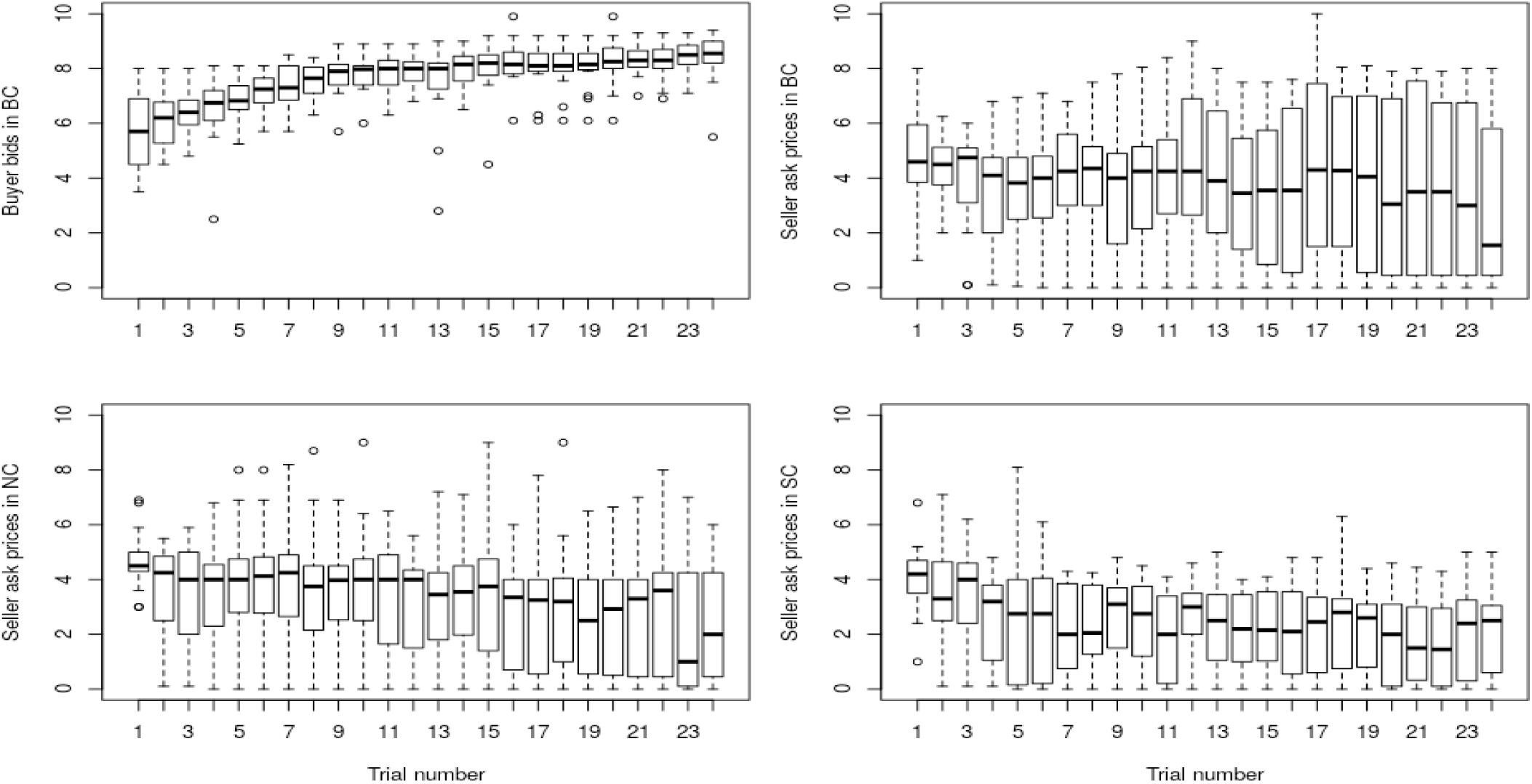
Opponents’ (buyers and sellers) prerecorded behavior in each market type. When the notches of two plots do not overlap this is strong evidence that the two medians differ (see Chambers et al. 1983; R Core Team, 2016).

The experimental session itself lasted for about 30 minutes, amounting to 45 minutes together with instructions and payments. After the task, subjects were rewarded according to the following reward scheme: a fixed compensation of 50RUB on account of participation, and in addition a bonus equal to the sum of the profit (in the task monetary units, MU) earned in three random trials multiplied by 5 RUB/MU. Therefore the reward was a number of RUB bounded between 50 and 200. Mean reward was 114 RUB (about 2 USD at the time of the experiment, when adjusted to PPP it was about 5$, which is roughly sufficient to cover the cost two lunches in the student canteen).

## 4. Learning models

Learning models in economic games have emerged from the study of empirical play in repeated games as a tool to study convergence to non-cooperative equilibria (Mookherjee & Sopher, 1994; Erev and Roth, 1998; 2014; Camerer & Ho, 1999; Feltovich, 2000; Sarin and Vahid, 1999, 2004; Sato et al., 2002; Grosskopf, 2003; Iwasaki et al., 2007). Perhaps the most popular of these is the “reinforcement learning model, in which players pay attention only to their own payoffs, and tend to repeat actions that have led to good payoffs in the past.” (Erev and Roth, 2014, p.10819). This approach emerged in natural sciences, but gained much popularity in economics literature because of its flexibility and descriptive capacities (Chen and Chen, 2009). Other adaptive learning models include deterministic (Brown, 1951) and stochastic fictitious play (Fudenberg and Kreps, 1993; Monderer et al., 1997), belief learning (Ioannou and Romero, 2014), along with their more involved alternatives and generalizations, such as *experience weighted attractions* (EWA – Camerer and Ho, 1999; Camerer et al., 2002). Although no model seems to be of one-size-fits-all type (Salmon, 2001; Mohlin, 2014; Fudenberg and Levine, 2016), reinforcement learning is the most popular in a range of applications, especially dealing with ‘passive’ learning in standard environments.

Following this approach, we contrast learning algorithms of reinforcement learning (RL) type, furnished with a function which estimates values of actions and states, and directional learning (DL) algorithms, lacking such function. These models impose rather mild requirements on human rationality: agents should not be able to observe strategies, but choices; learning should be based on a parsimonious algorithm; and unobservable parameters for the opponents are not considered.

Reinforcement learning (RL) is a strategy for maximizing the profit of an agent placed in an uncertain environment by means of the storage and updating of a value function which encompasses all necessary information to act in an optimal manner (Sutton & Barto, 1998). In the simplest, model-free form, when the environment can be modeled as a *Markov decision process*, the optimal strategy can be found by value iteration of the Bellman equation, which involves systematically sweeping the whole state space. However, this is unfeasible or unpractical for most situations, so most applications of the RL algorithm use the dynamic action-value updating equation of the RL model whose generic form can be written as

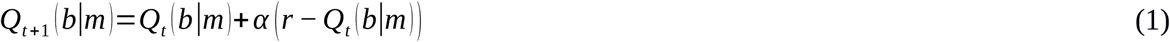

With the interpretations for our case, Q(m | b) is the action-value function with a value for each bid *b* from the set of possible bids *B*, conditional upon market type *m* at trial *t*, and α is the learning rate regulating the speed of action value updating. For comparison, (1-*α*) corresponds to Roth & Erev’s (1995) parameter ε or “persistent local experimentation” (“gradual forgetting” *φ* and “cutoff” μ parameters are both null constants here). Action values were learned independently for each of the three market types.

The policy for selecting a bid in each trial for the most successful model is a logit (softmax) function:

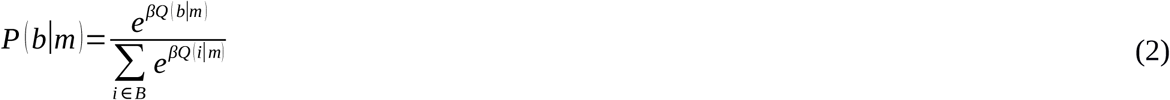

where P(b | m) is the probability of choosing bid *b* in market type *m*, and *βQ* is the elasticity parameter which captures the importance of the reinforcement signal, *B* is the space of actions (bids), which in this study contains 101 elements. Roth & Erev (1995) normalized *Q(b*|*m)* values simply dividing by their sum because their *Q(b*|*m)* values were defined to be always positive.

Many adaptive models emerge from this simple scheme, corresponding to alternative learning rules. In our case the problem is greatly simplified because there is essentially only one state (per market type), hence our problem lies exclusively in choosing a single-dimensional value – the preferred action (bid) (Lee et al., 2004). However, unlike most RL problems in natural sciences, in our game-theoretic context players could adopt learning strategies based on “rational” pre-game analysis or on the behavior of the other players (Lee et al., 2005). In order to address these possibilities, we consider a broader state and strategies space, and learning models which account for the relevant prior information about the structure of the decision problem.

### 4.1 Model-based RL with counterfactual action-values updates

A first step in application of the general RL model is to take advantage of the fact that action values are ordered. One option is the similarity-based ‘‘spillover’’ of payoffs from a chosen bid to neighboring bids explored by Sarin and Vahid (2004). A more informed strategy would be to control the spillover by using information about foregone payoffs through counterfactual learning. This strategy is carried into effect explicitly in those model-based RL models which asymmetrically update with every choice bid, conditional on both the bid value and the feedback. Counterfactual learning is also carried into effect, although indirectly, in DL algorithms, which incorporate knowledge about the structure of auction games by adopting the apparently simplistic but informed strategy of ‘nudging’ up or down the preferred bid value in congruence with the feedback direction.

Counterfactual learning is an extension of model-free RL where the value function is updated contingent not only on the currently chosen action feedback, but also on non-chosen actions based on a model about the contingent rewards of foregone actions. In such cases, value updating occurs for actions which were not chosen, but which are nevertheless updated based on the assumption that they would have been updated had they have been chosen. Counterfactual learning is implemented according to the following rule: for every bid b selected, if it is accepted (rejected), increase (decrease) the value of the action-value function for all *a>b*. An open question is how much to decrease or increase the value, and for which actions. One possibility is to update values conditioned on the outcome of the current transaction, and do it asymmetrically respect to the current bid (action). This yields four possible combinations of accepted/reject bid and bid values higher/lower than the currently bid:

If accept:

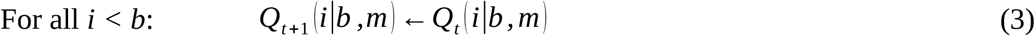

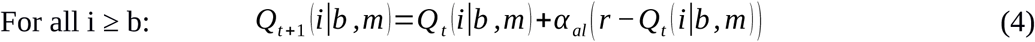

If reject:

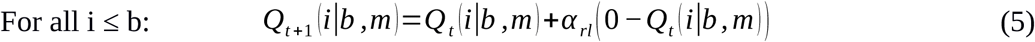

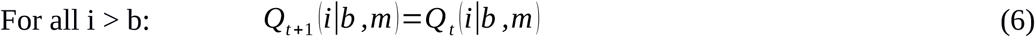

where α is the learning rate, and *r*_*i*_ is the counterfactual reward, that is, the reward the player would have received had she selected the bid *i*. For the current trial bid *b, r*_*i*_ = *r*_*b*_ = *r*, the reward actually obtained.

### 4.2 Directional learning

DL is a more simplified adaptive strategy which, crucially, assumes that the available actions are ordered under some consistent relation, as in the case of a well-ordered set of available bids. The term DL was coined by Reinhard Selten (Selten and Buchta, 1994) to describe a “learning direction theory” applicable to tasks involving the uncertain dynamic evolution of a one-dimensional quantities. An agent acting in accordance with DL will exhibit a simple Markovian dependence on the immediately previous feedback, and dispense with any kind of value function, unlike RL. Adaptations of RL using this idea have been proposed by Gullapalli (1990) and van Hasselt & Wiering (2007). DL is suited to model the typical round-to-round behavior in bargaining tasks where proposers tend to increase their offers following a rejection and reduce it after being rejected (Mitzkewitz and Nagel, 1993), although this has been contested (Roth and Erev, 1995). Research analyzing the relationship between DL and RL exists also in a bargaining task where a combination of both RL and DL is endorsed by Grosskopf (2003).

DL is effectively a policy (a rule for selecting actions) which operates without the need of action value functions, in contrast with RL policies, which hinge on the distribution of value functions to yield probabilities for choosing actions. Nonetheless, DL posits some degree of model-based adaptive learning by making use of an error signal. However, unlike the scalar reward prediction error of RL, this “directional signature” is binary, because its only purpose is inducing a sign in a one-dimensional span of choices.

DL has been found to describe accurately a large body of experimental data (Mookherjee & Sopher, 1997; Erev & Rapoport, 1998; Erev & Roth, 1998a; Slonim & Roth, 1998; Camerer & Ho, 1999, Martinez-Saito et al., 2019), but it does not furnish testable quantitative predictions: the magnitude of this adjustment remains unspecified. This indeterminacy has hindered the development of theories of behavior in the spirit of DL, in search for parsimonious models that do not require the incorporation of ad hoc parameters.

We implemented DL algorithms by, in every trial *t* of market type *m*, picking a bid from a unimodal probability distribution *P(b*|*m)* centered in the preferred current bid (PB), which we denote by *A*. If the selected bid is accepted (rejected), then the PB is increased (decreased), determining the PB for the next trial according to the equation:

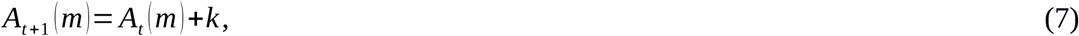

where *A*_*t*_*(m)* is PB for trial t in market type *m, k* is the gain, i.e. the PB adjustment, typically conditioned on the specific algorithm used, as detailed below in each agent model type.

As in the previous case (model-based RL with counterfactual updating), this leaves unspecified how much to decrease or increase the PB. DL has been found to describe accurately a large body of experimental data, but it does not furnish testable quantitative predictions, since it does not prescribe how to adjust actions upon miss other than the *direction* in which the actions should be adjusted. In this study, we have created and fitted several adaptive learning models based on DL. The numerous schemes we tried on the dataset are succinctly described next:

*Naive DL*. This is the basic model, which we did not implemented in practice, but it is the basis of all other DL models used in the current study. It consists in simply ‘nudging’ the bid up and down by a fixed amount *n*, contingent on the outcome of the transaction:

The naive learning model stipulates adjustment from the previous trial is a function of success or failure only, and is being made at a constant rate:

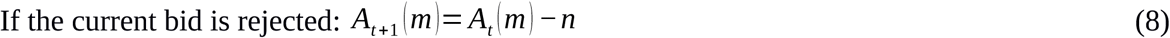

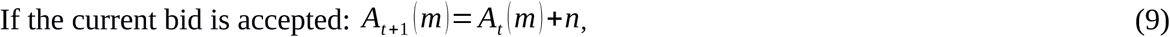

where *n* is a constant to be estimated. The policy is defined by simply picking PB in the next trial.

*Naive DL with Gaussian noise*. It behaves like Naive DL, with the difference that the policy accounts for noisy decision making with a Gaussian distribution function of bids around the PB:

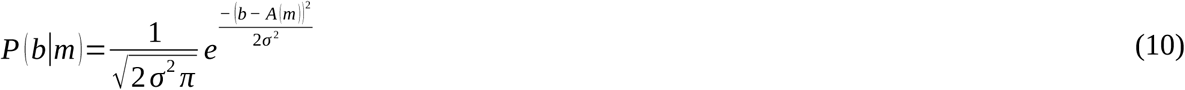

where *σ* is the standard deviation and *A(m)*, which is equal to the PB for market type *m*, is the mean.

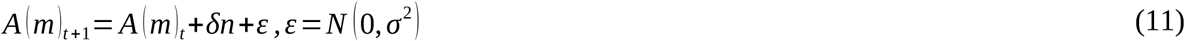

This model is identical to random walk with drift driven by acceptance or rejection of the previous bids. In this model we estimate parameters n and σ.

*Naive DL with asymmetric leptokurtic (Laplacian) noise*. Its bid distribution of bids is asymmetric and non-gaussian, specifically with fatter tails and thinner peak (more kurtotic, or leptokurtic) than the Gaussian distribution (Figure S1, Figure 1C). Thus, in order to improve the fit, we used an asymmetric leptokurtic distribution to accommodate this behavioral trait.

For *b > A(m)* after previous trial rejection

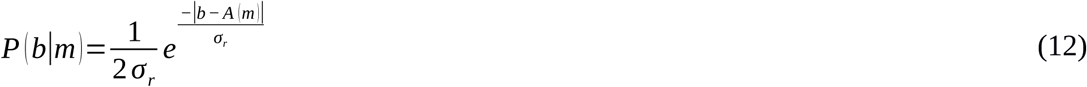

for *b < A(m)* after previous trial acceptance

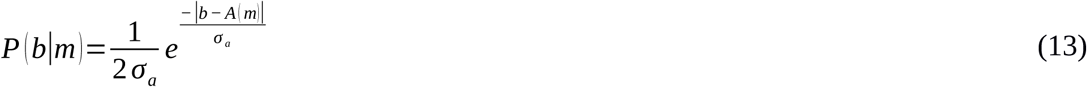

and for the rest of (rare) cases,

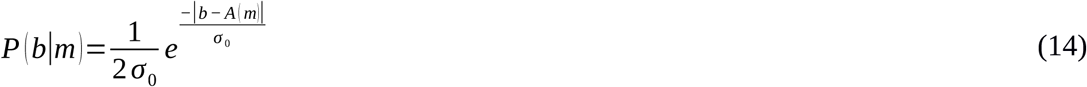

where *P(b*|*m)* is the Laplace distribution of bids *b* for market type *m*, and *σ*_*r*_, *σ*_*a*_, *σ*_*0*_ are parameters proportional to the standard deviation of the Laplace distribution. This captures that intuition that the tail above PB after rejections is fatter than the tail below PB after acceptances.

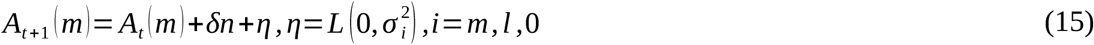

This model is identical to random walk with drift driven by acceptance-rejection of previous bids and leptokurtic noise. Here we estimate n and *σ*_*r*_, *σ*_*a*_, *σ*_*0*_.

*DL delta rule with Gaussian noise*. Another possibility is to update values conditioned on the outcome of the current transaction, and do it asymmetrically respect to the two sides of PB. This can be done either fixing different gains for increasing or rejecting PB, or by conditioning the gain on the value of the current PB and the reward received. This model incorporates reward prediction, as in RL, the difference being that it is conditional not upon updated action values, but upon updated PB:

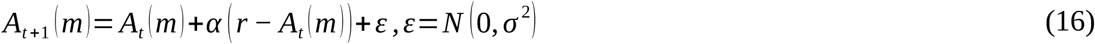

where *α* is the effect of gain, akin to the learning rate in RL, referred as to *delta rule* in this context. The policy for bid selection is again a Gaussian distribution centered on PB. Here (r is reward of the previous period), and we estimate the learning rate *α* and *σ*^2^.

*DL delta rule with asymmetric leptokurtic noise*. This model combines both the asymmetric leptokurtic policy distribution and the delta rule-based updating of PB:

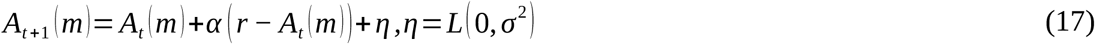

This model turned out to be the most successful empirically (see Section 5). In order to estimate the exploration-exploitation trade-off, we introduced an additional parameter (*ρ*) for the proportion of trials with explorative (risky) versus exploitative (safe) bids. The best fitting model had 5 parameters (*α* = 0.59, *σ*_*a*_ = 0.45, *σ*_*r*_ = 0.60, *σ*_*0*_ = 0.43, *ρ* = 0.36).

### 4.3 Belief learning of opponent choices

Finally, we modeled learning with an algorithm which estimates opponent’s behavior with Gaussian kernel density estimators, the bid density estimator learner (BDEL). The term “belief learning” originally refers to a interactive play strategy where players’ actions are based on the selection of choices corresponding to the weights assigned and updated according to the past behavior of other players (Goeree and Holt, 1999). Such beliefs are typically formed according to weighted fictitious play and used to calculate attractions values to each strategy, including those which have not been chosen in that period (Camerer and Ho, 1999; Gigerenzer and Selten, 2002; Lee et al., 2005). This density is estimated with a Gaussian kernel fed with the feedback from past trials. In T1 and T2 treatments, belief learning plays a main role only in BC market because of limited feedback, hence in these models we assume that the opponent’s choice was uniformly distributed in the support of the bid line congruent with the feedback (i.e., if in NC market the player bid was 6 and it was rejected, the player assumed that the opponent seller chose an ask price with uniform probability in the segment [6.1, 10], etc.). In T3 treatment, beliefs about both sellers and competitor buyers can be updated every trial.

On every trial, the player is assumed to calculate the utility function (U) of choosing a bid by weighing, for every bid value, the probability that the opponents chose their respective bids and/or ask prices based on the density estimated from the Gaussian kernel density estimators, thereby estimating the probability that the bid was accepted, and finally multiplying this probability by the profit (10 minus their bid) endowed in case the bid was accepted.

Opponents’ choice densities were estimated independently for each market type. As for RL models, the policy for selecting a bid in each trial for the most successful model was a conventional logistic function:

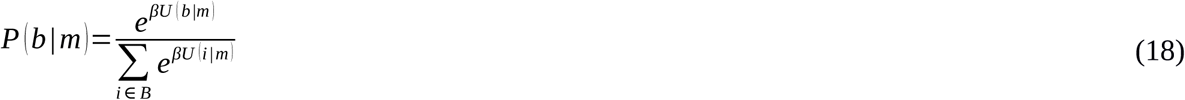

where *P(b*|*m)* is the probability of choosing bid *b* in market type *m*, and *βQ* is the elasticity parameter with respect to estimated utilities of the opponents in the space of actions B.

The above models were estimated for each market type, with the simplified state space. Natural action space consists of 101 actions, to wit, the sequence from 0 to 10 in steps of 1/10. Note that although the support of this space support is discrete, neural mechanisms based on continuous or coarser representations are also plausible. For the purpose of scouring for the best fitting model, we consider also a coarser action space of just 11 points (from 0 to 10 in steps of 1). This choice was made for technical reasons: convergence of RL models is remarkably difficult for games where there is a large number of states or actions. Therefore, we either simplified the initial action-values using a three-parameter (as opposed to 101) beta distribution, or simply used the first round bids as initial conditions, saving all 101 parameters for the actually meaningful parameters, i.e. learning rate and elasticity.

### 4.4. Hypotheses

In the light of the above discussion, our first hypothesis (H1) is that behaviour in full information (T3) shall result in more efficient learning than in partial information (T1) and scrambled information (T2) treatments. Further, we expect (H2) that structural uncertainty in treatment T2 exposes subjects to true uncertainty, and shall result in larger efficientcy losses than strategic uncertainty in treatment T1 relatively to full information treatment T3. Finally, (H3), inasmuch as information acquisition is difficult and challenging, we expect that subjects will use simpler learning rules in most loaded environments, such as T2 relatively to T3.

## 5. Results

### 5.1 General

First, we compare the bids of the first trial of each treatment. This decision provides an estimate of a combination of subjects’ beliefs about opponents’ player types and a pre-game analysis of each market type. Except for SC and NC market types in T1 treatment, first trial bids were significantly different among market types, which is especially obvious in the full information, T3 treatment (Figure 3). This confirms that subjects understood and factored in the structural uncertainty, even in case of incomplete information (T2 treatment), although to a lesser extent in this last case.

**Figure 3.**
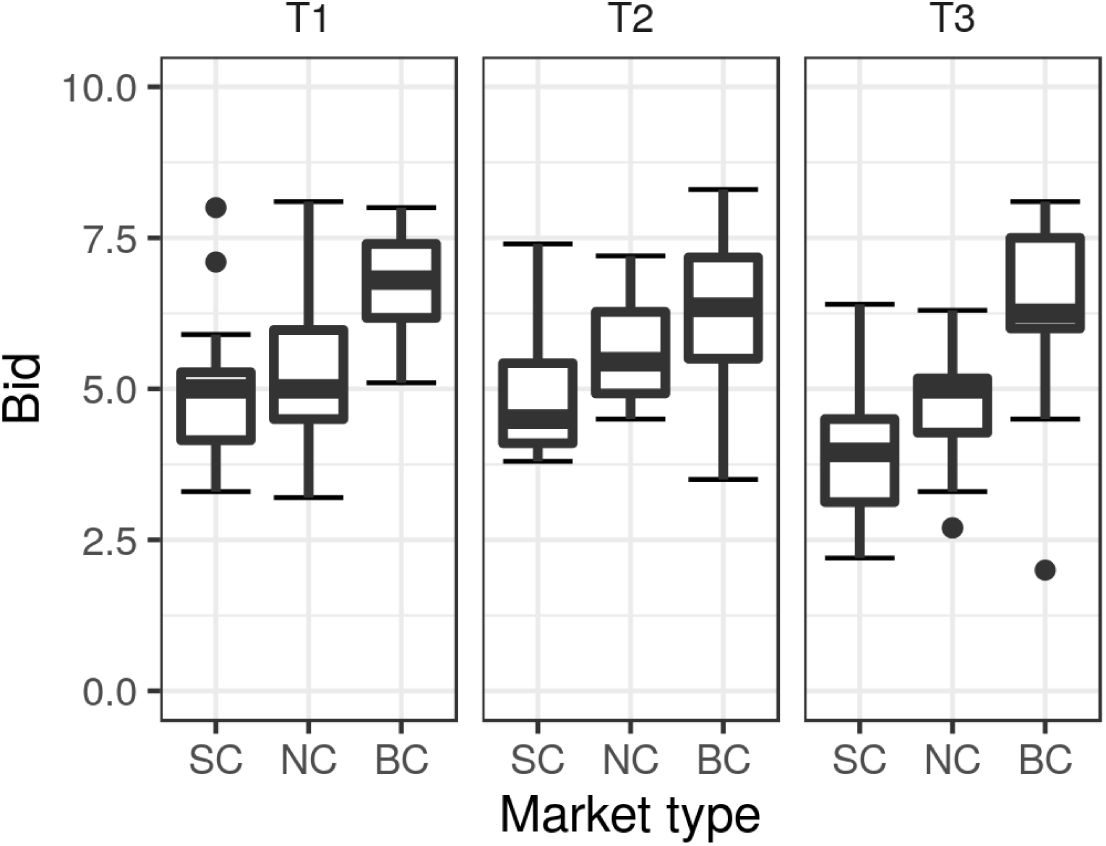
First bids across treatments (experiments) and market types. In all treatments, one-way ANOVA rejected the null hypothesis that first bids were indistinguishable different across market types: T1 (p < 10e-5), T2 (p < 10e-3), and T3 (p < 10e-5).

We now turn to overall results. General statistics of trades is provided in Table 2 and illustrated further on Figures 4 and 5.

**Figure 4.**
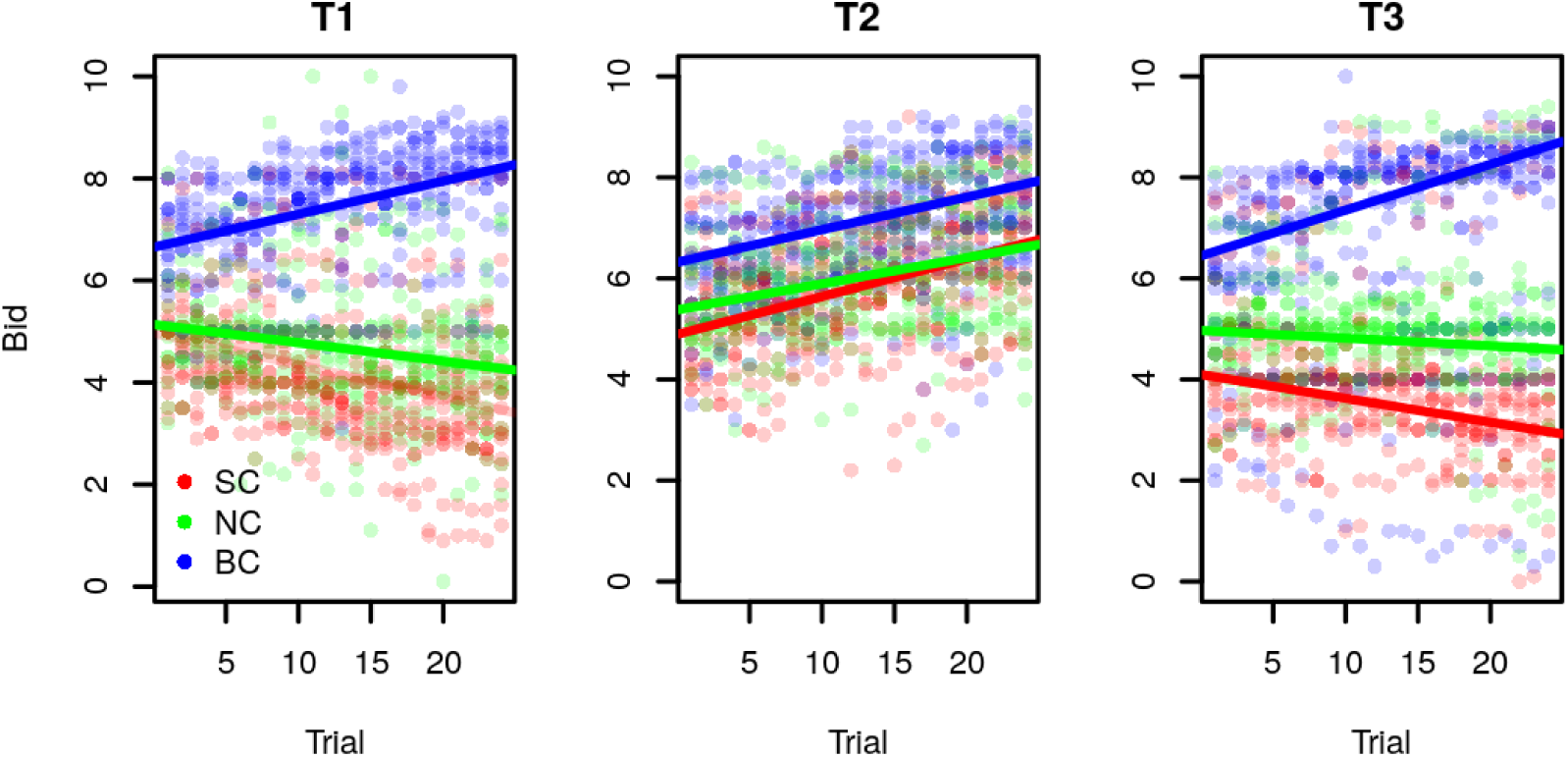
Bid evolution across treatments. Left (T1 treatment): Slope coefficients of the regression line are: -0.06 for SC (p < 10e-10); -0.04 for NC (p < 10e-5); and 0.06 for BC (p < 10e-15). Middle (T2 treatment): Slope coefficients are: 0.08 for SC (p < 10e-15); 0.05 for NC (p < 10e-11); and 0.06 for BC (p < 10e-15). Right (T3 treatment): Slope coefficients are: -0.05 for SC (p < 10e-9); -0.02 for NC (p = 0.02); and 0.09 for BC (p < 10e-15).

**Figure 5.**
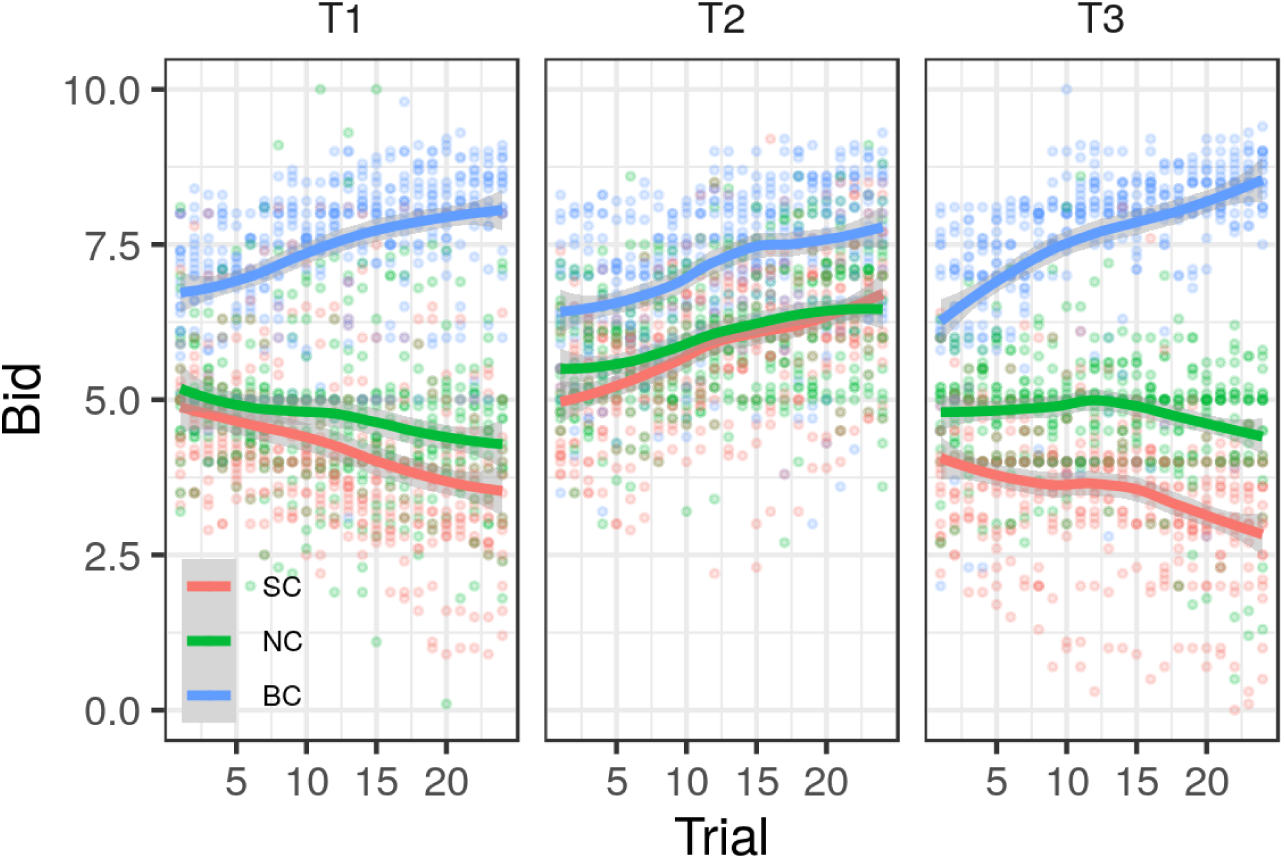
Bid evolution trends for all treatments and market types. Local quadratic polynomial regression curves with 95% confidence intervals plotted.

Overall, 71.8% (2714/3888) of transactions were successful. Breakdown table of successful transactions by market type and experimental treatments T1, T2, and T3, ratios varied significantly, with typically lower success rates in T2 treatment, except for BC market type (Table 2). Differences between success rates in this and the other two treatments are all highly statistically significant, which speaks in favour of our H2, confirming the expectation that structural uncertainty will result in larger efficiency losses than strategic one.

### 5.2 Evolution of bids

The dynamics of learning in all markets can be characterized by means of linear regression against an intercept and number of bids (Figure 4) and local quadratic polynomial regression curves fit across the 24 blocks (Figure 5). Several things are worth notice. First, the evolution of bids in T1 reveals gradual but less than full convergence to the Bertrand-Nash equilibria in BC, and a downward trend for SC, again corresponding to the equilibrium component of the Bertrand type, but to a lesser extent (Figure 4). Convergence in these market types is not complete, which may be attributable to short learning series. Bids under NC are indistinguishable from equal split strategy, consistent with the Nash bargaining solution in case of equal bargaining power.

Second, in T1 and T3 treatments learning is steady, but not complete in the predicted directions in both BC and SC market types, with all slopes being significantly different from zero. By contrast, bids in T2 treatment exhibited an upward evolution for all market types, with the difference between the learning slope being insignificant across treatments, but higher intercept in SC in comparison to BC and NC market types. This can be explained by risk aversion in the scrambled treatment: subjects are afraid of not meeting the bid of the (relevant) seller, and raise their own in order not to miss the chance to make a positive, albeit smaller, profit. Appendix C provides a simple theoretical argument for this conjecture.

Finally, it is worth notice that learning in SC is less steep than in BC, and closer to the equal split prediction of NC than to the Bertrand prediction. This can be attributable to risk attitudes: as described above, transaction partners were determined by random matching, hence the corresponding higher risk of thwarting the deal also affecting the direction of bid adjustments.

These findings can be summarized as follows:

#### Result 1

Convergence of bids in the direction of Bertrand Nash equilibria takes place for both BC and SC, and the effect is stronger in BC than in SC market types.

#### Result 2

Bids in T2 treatment are systematically increasing, which can be explained by risk aversion of the bidders who are more likely to fail the deal in the scrambled market.

Our primary explanation for the difference across markets in Result 1 is risk attitudes. Subjects who know that they may be misled by the signal compare the expected loss from making higher offer and chances of not making the transaction. Since there are at most two players, the probability that the transaction will fail looms larger, which forces them to increase their bid in all cases. Appendix C makes the point more precise.

Two further tests can bolster the argument of Result 2. Figure 6 shows the mean differences of bids between SC and NC (SC-NC), and BC and NC market types (BC-NC) separately for T1 and T2 treatments. In T2, SC and NC bids become indistinguishable, while BC bids are higher than NC bids by a constant offset of about 1 unit, with rather small variance (see Figure 6, right). By contrast, in T1 (figure 6, left), the variance of BC-NC is much larger, with the offset increasing on average to 2.8.

**Figure 6.**
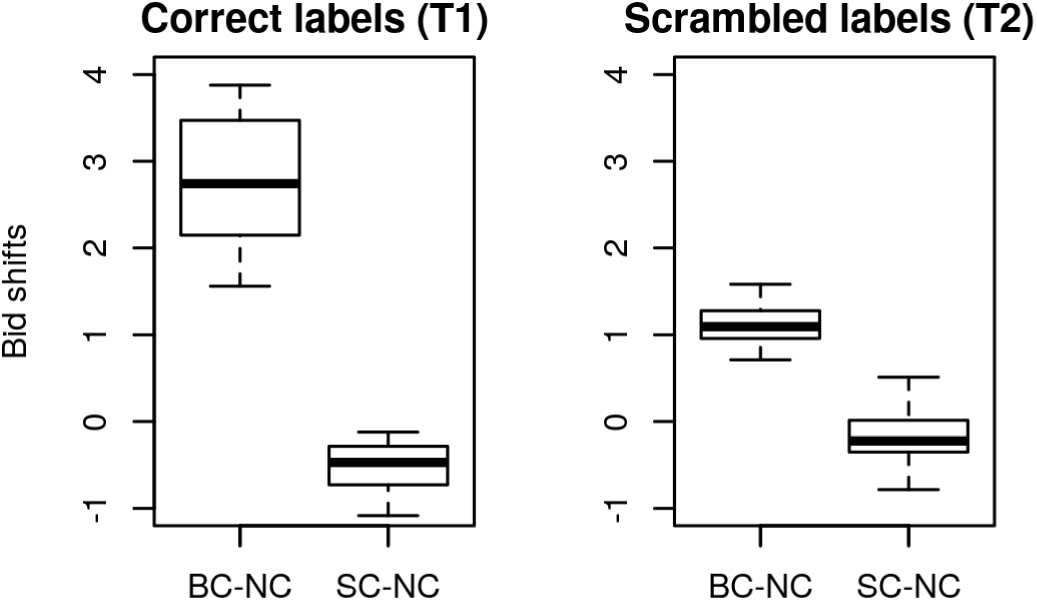
Between subjects stage-wise mean bids of SC and BC respect to NC for T1 (left) and T2 (right). Box hinges are the first and third quartiles. Whiskers extend to the most extreme data point which is no more than 1.5 times the interquartile range from the box.

As another robustness check, we look at homogeneity in subject’s bidding patterns. In order to rule out the possibility that (across subjects) performance was determined by players’ heterogeneity, we explored the successful transaction rates by averaging rates across subjects (Figure 7). The distribution of acceptance rates is unimodal for all three treatments, which implies that individual variations about the mean are not systematically affected by the treatments.

**Figure 7.**
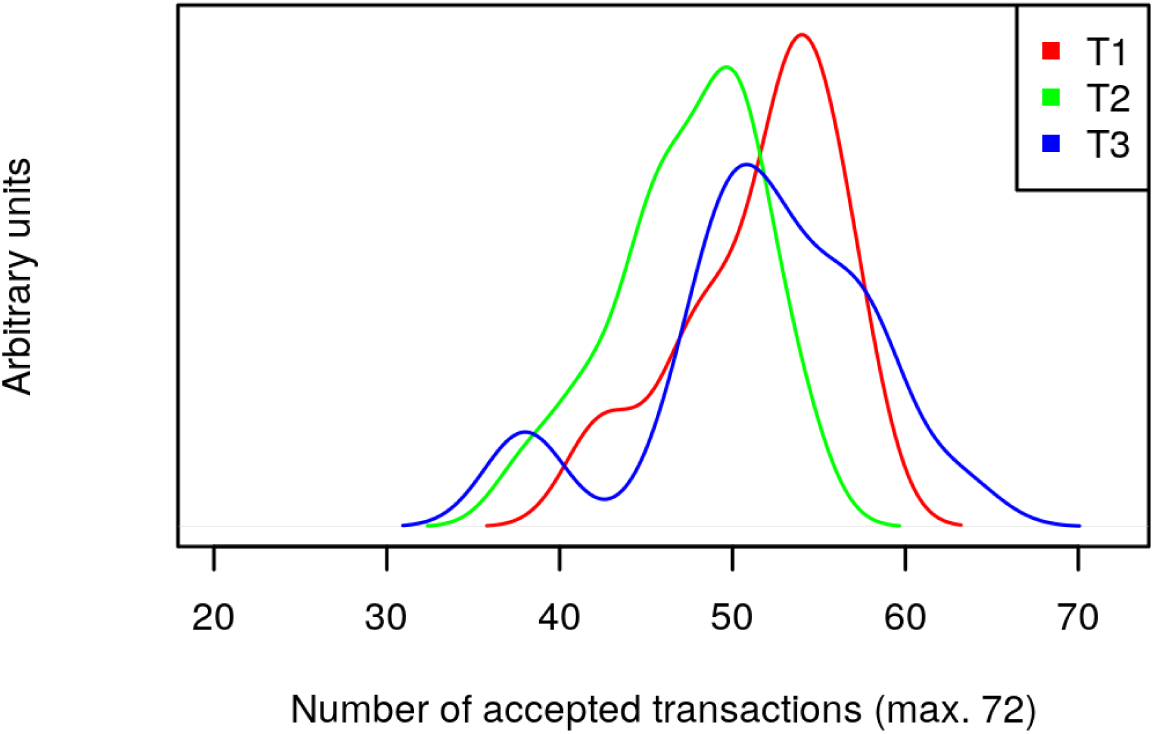
Density estimates of percentage of accepted transactions across subjects. A Gaussian kernel was used for smoothing (*density* function in R package *stats* with default parameters, R Core Team, 2016). The bandwidth of a Gaussian kernel density estimator is 0.9 times the minimum of the standard deviation and the interquartile range divided by 1.34 times the sample size to the negative one-fifth power.

### 5.3 Between-treatment differences

Let us now examine the influence of the treatments on the average profit earned. As expected, subjects earned significantly more in T3 than in T1 treatment, and in T1 than in T2 treatment (Figure 8). So, across experiments, T2 trials profit was qualitatively different and lower than the profit of T1 and T3 (uncorrected one-sided Mann-Whitney U tests for between-subjects mean profits across the three treatments: T1-T2: W = 304, p-value = 3.777e-06; T3-T2: W = 324, p-value = 1.611e-07; T3-T1: W = 172, p-value = 0.3819). This confirms our hypothesis H2: in a given market environment, structural uncertainty has more dramatic consequeces than strategic one.

**Figure 8.**
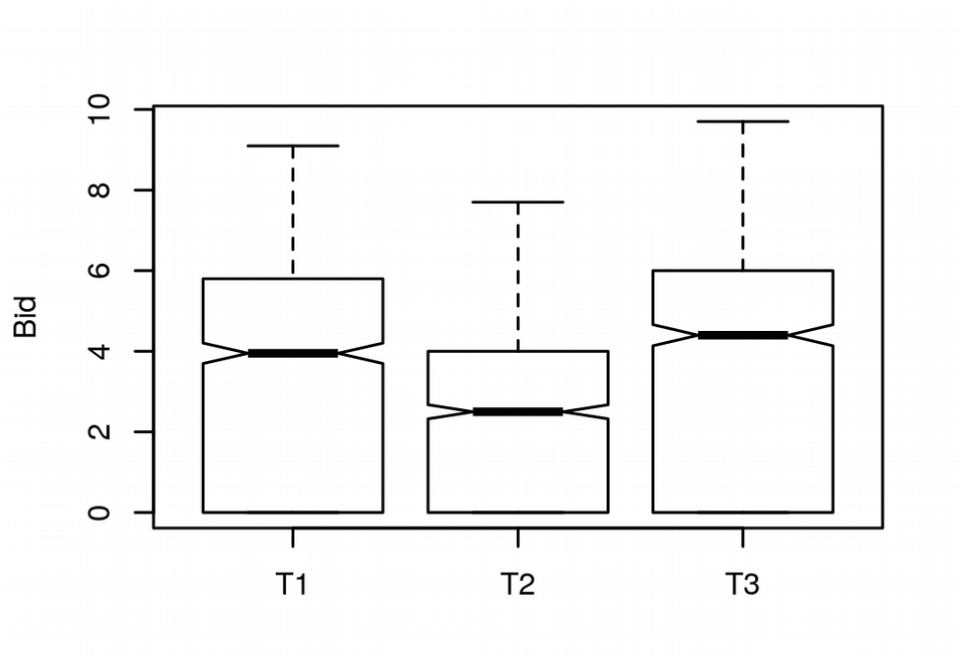
Profit statistics. Earned average profit in each treatment (experiment) across all trials.

#### Result 3

Profit under T2 treatment is systematically lower than under T1 and T3 treatments, while the last two are not systematically different.

### 5.4 Trial-by-trial dynamics

Now we turn to examine the trial-by-trial dynamics of bidding behavior. The most conspicuous pattern of consecutive bid adjustments within each market type is that their distribution is leptokurtic (with a central peak more pointy and longer tails than those of a Gaussian distribution) and skewed (asymmetric). This pattern recurred in all sessions. Current bid distributions conditioned on an acceptance in the previous trial of the same market type were skewed towards the left (towards lower bids), whereas when conditioned on rejection the skewness sign was opposite (towards higher bids). Moreover, distribution variances were consistently larger in case of previous trial rejection (Figure 9, Table 3). This allows us to formulate our next result:

**Table 3.** Statistics of bid adjustments between trials of the same market type. Excess kurtosis is defined as kurtosis minus 3.

| Treatment Statistics | T1 | T2 | T3 | Total aggregated |
| --- | --- | --- | --- | --- |
| Acc Median | -0.1 | 0 | 0 | 0 |
| Acc Mean | -0.15 | -0.13 | -0.18 | -0.16 |
| Acc Standard deviation | 0.79 | 0.86 | 0.75 | 0.80 |
| Acc Skewness | -0.97 | -0.36 | -2.08 | -1.04 |
| Acc Excess Kurtosis | 11.48 | 4.20 | 12.37 | 8.84 |
| Rej Median | 0.3 | 0.3 | 0.4 | 0.3 |
| Rej Mean | 0.42 | 0.41 | 0.53 | 0.45 |
| Rej Standard deviation | 0.92 | 1.01 | 1.11 | 1.02 |
| Rej Skewness | 1.40 | 0.28 | 0.88 | 0.80 |
| Rej Excess Kurtosis | 8.89 | 3.94 | 9.97 | 7.83 |

**Figure 9.**
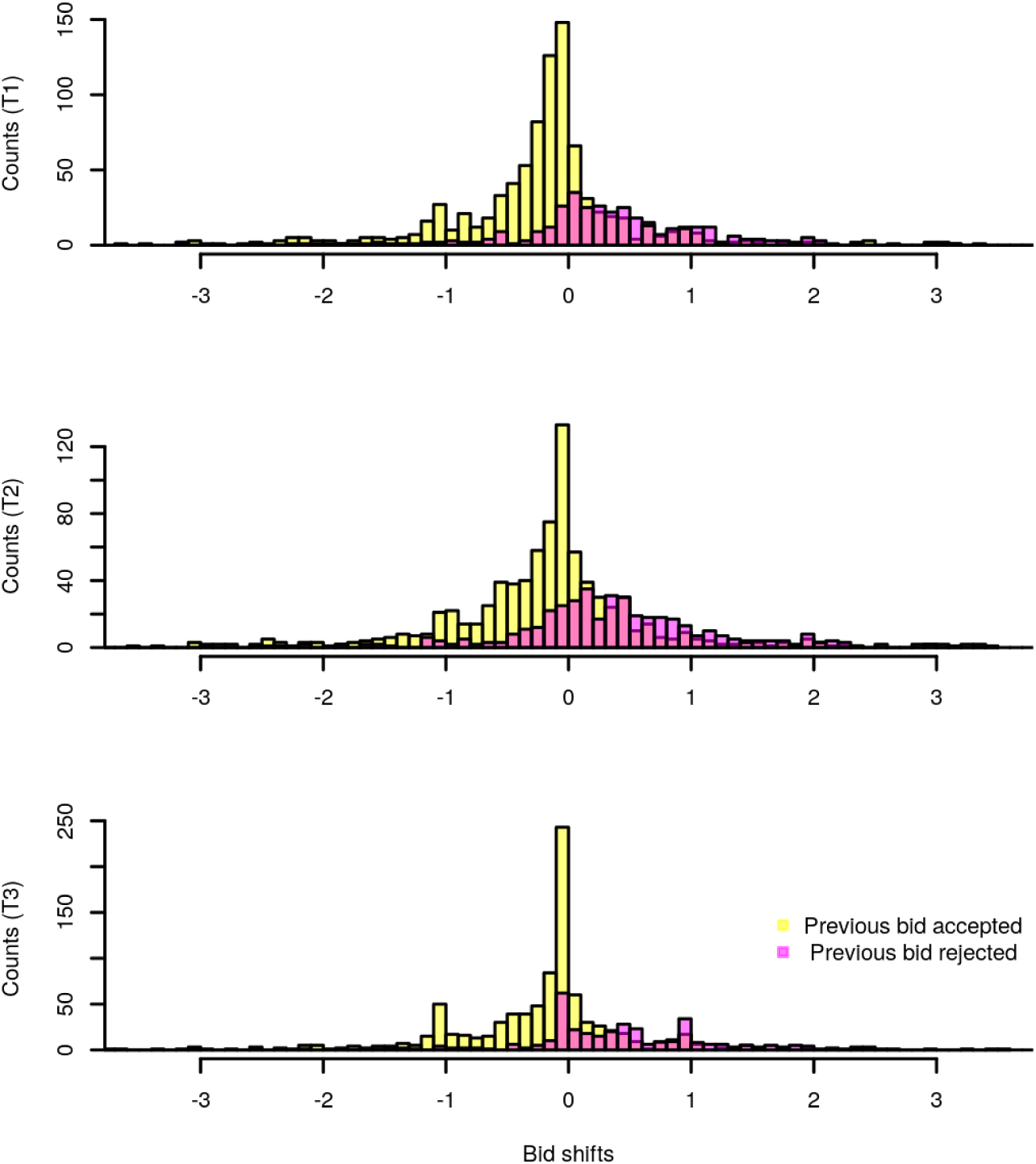
Bid choice shifts conditioned on the previous outcome across treatments.

#### Result 4

Bids following successful transactions are typically lower, and bids following rejected transaction – typically higher than in the previous period.

### 5.5 Learning models

Finally, let us come to estimations of the learning models, which are summarized in Table 4. Model fits, as indicated by BIC scores, allowed to discriminate plainly between Null (benchmark model with only 1 parameter fitting the mean), RL, BDEL, and DL strategies (Figure 10), and indicates that DL as a clear champion, confirming our hypothesis H3. Uniform BIC scores across treatments suggests that subjects used in essence the same strategy for all T1, T2 and T3 treatments, plausibly a DL algorithm. Aggregated BIC scores for 1296 trials were 1037.2, 1234.8, and 1071.2 for T1, T2, and T3 treatment respectively. Corresponding negative log-likelihood in nats per trial were 0.39, 0.46, and 0.41. RL algorithms fit was not only poor, but also lacked generative sufficiency (Figure 11), in agreement with Roth & Erev (1995). DL algorithms exhibited generative sufficiency in the sense that simulated artificial bidders pit against the same pre-recorded dataset as humans generated behavior resembling that of human subjects: the simulated learning curves of agents enacted by DL algorithms playing against the prerecorded dataset were qualitatively in close agreement to human behavioral learning curves (Figure 11). By contrast, learning curves of agents enacted by RL algorithms hardly exhibited any learning (not shown).

**Table 4.**
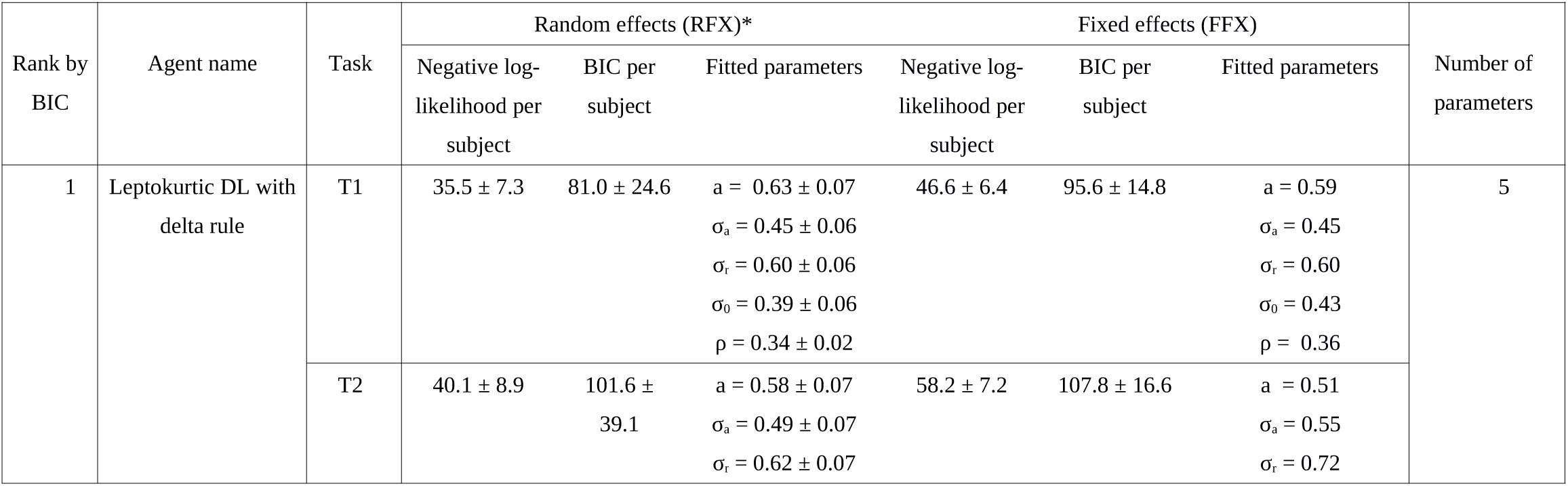
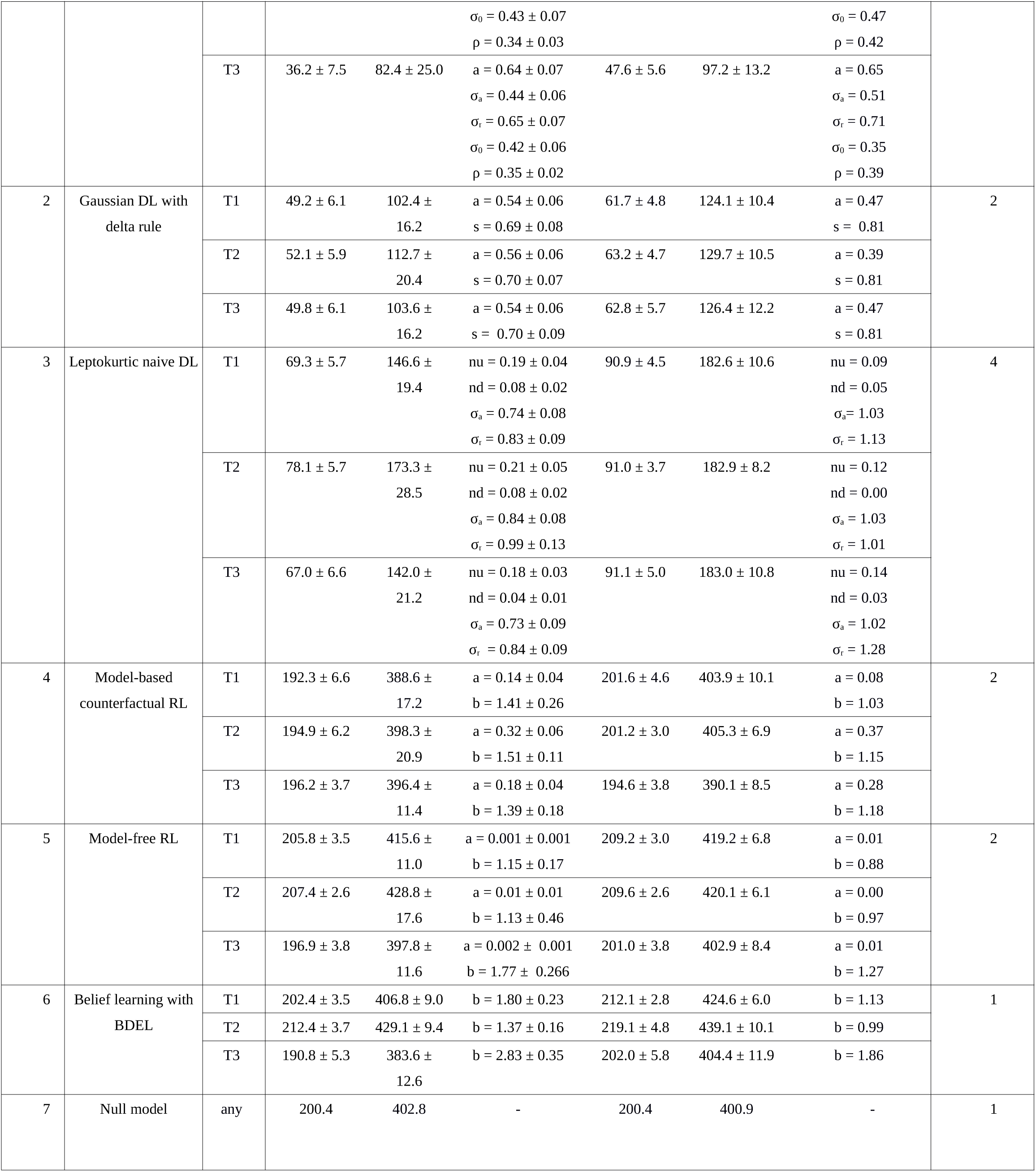
Ranks and BIC scores for all fitted algorithms. The sign ± denotes standard error of the mean (in FFX, individual BICs can vary across subjects even with yoked parameters). *Many instances of the random effects log-likelihood optimization did not converge. a: learning rate; b: inverse temperature; σ_a_, σ_r_, σ_0_ : variance of Laplace distributions; ρ: proportion of trials with explorative (risky) versus exploitative (safe) bids; up, down: fixed nudge size in the naive nudger algorithm.

**Figure 10.**
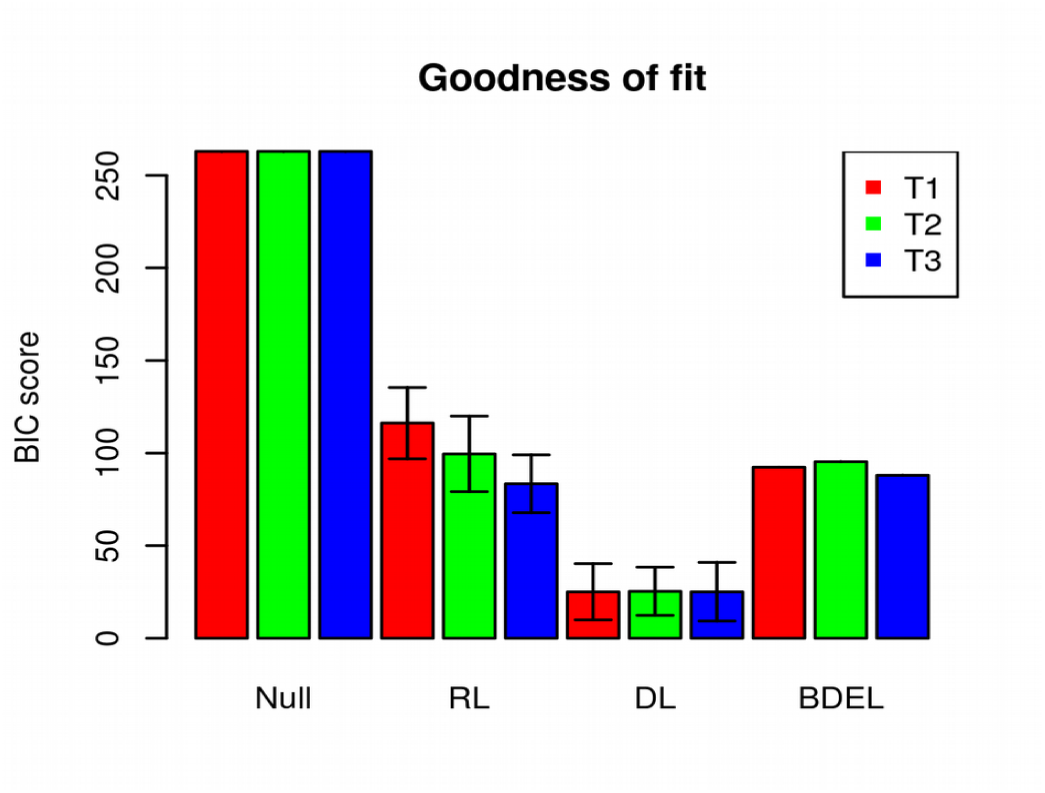
BIC scores of Null, RL, DL, and BDEL models and averaged across subjects. As described in section 4, we fit the parameters of each model yoked across subjects because the number of trials per subject (72) was comparatively small given the number of subjects. Error bars indicate 95% confidence intervals.

**Figure 11.**
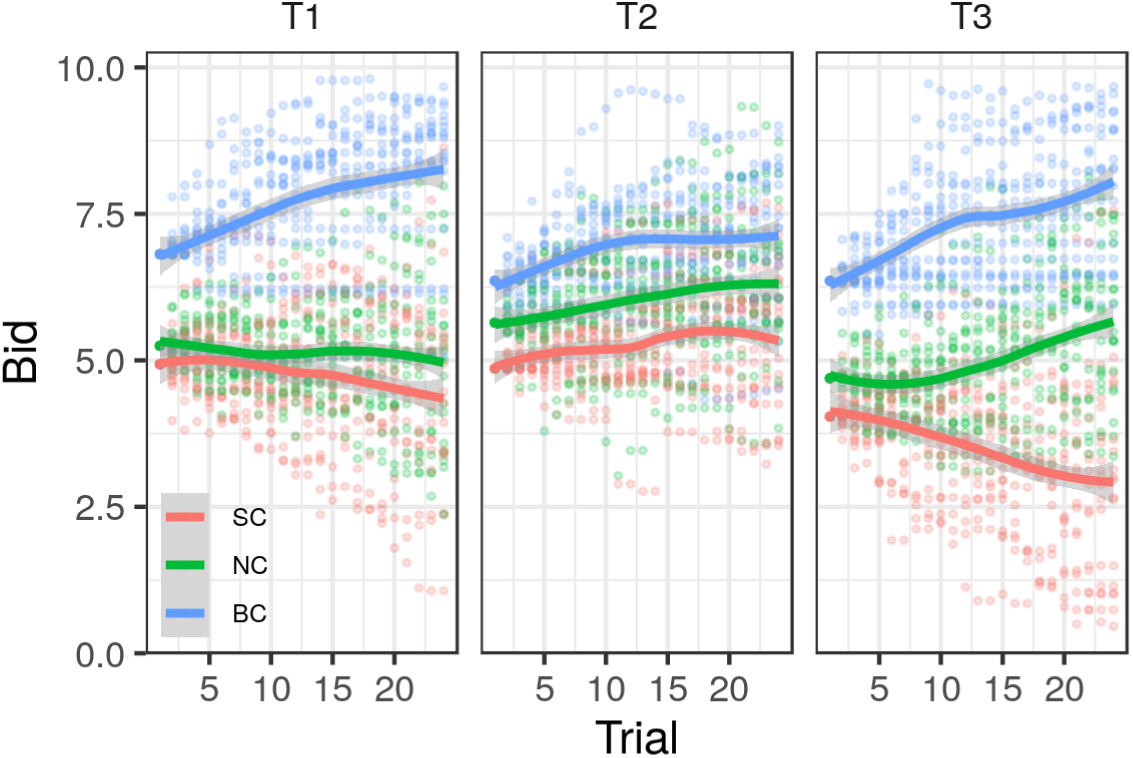
Learning curves of DL algorithms simulating artificial bidding agents pitted against the same prerecorded dataset subjects played against. Qualitative patterns of human subjects are reproduced (cf. Figure 5).

**Figure 12.**
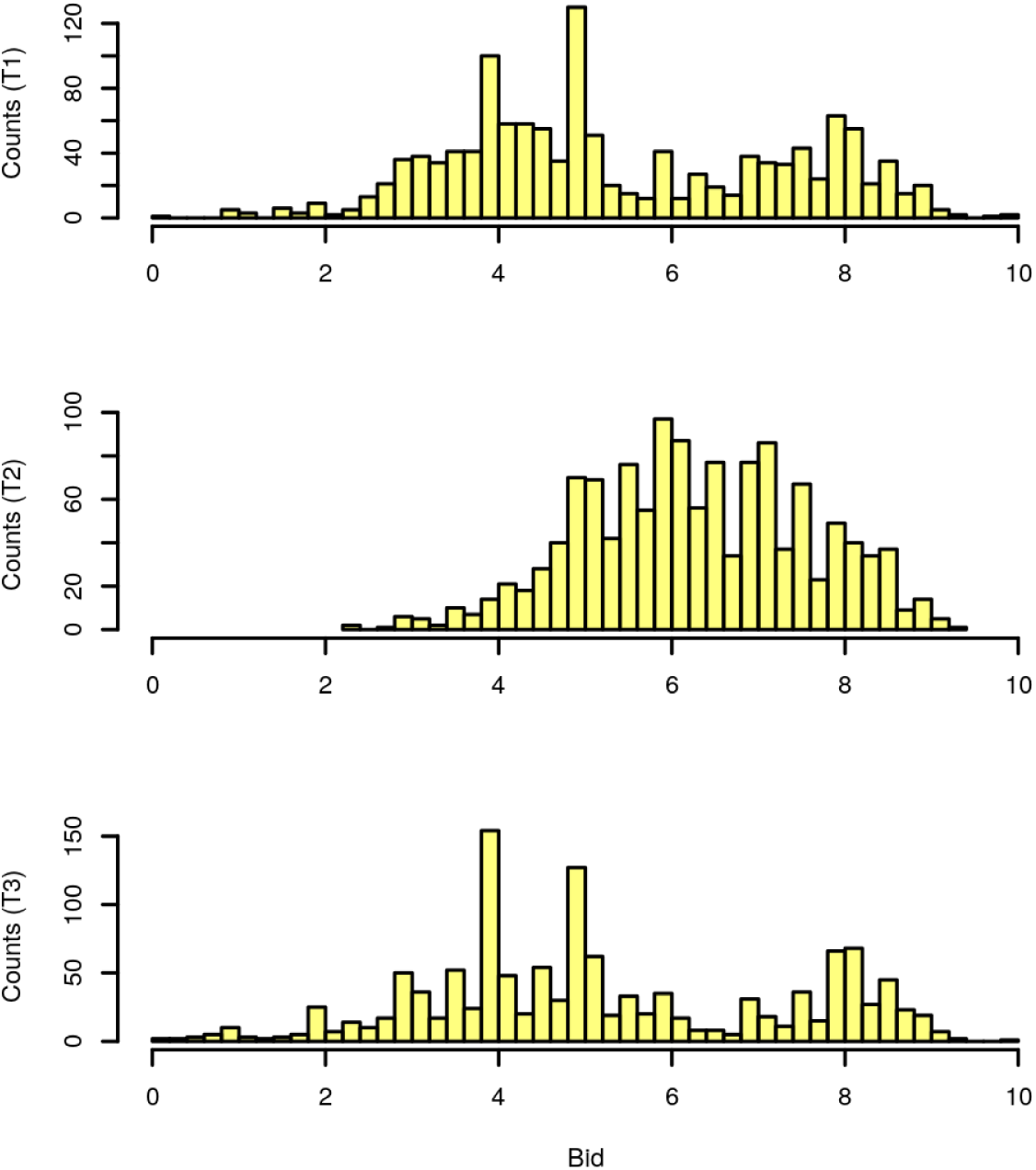
Distribution of buyers’ bids depending on the treatments. Finally, our empirical results convincingly show that individual buyers reveal risk aversion when facing potentially scrambled signals. Figure 12 shows the distributions of buyers’ bids in the three information conditions. Intuitively, it seems obvious that partial (T1) and full (T3) information conditions result in quite similar bidding patterns, which are not significantly different (Wilcoxon-Mann-Whitney p-value <0.1067). By contrast, bids under scrambled information are much larger than those under partial and full information (WMW p<2.2e-16 in both conditions). This evidence clearly suggests that buyers’ reaction to larger **ambiguity** about the true structural uncertainty exerts major influence on individual strategies, which are of larger importance than the strategic ones.

## 6. Discussion

In line with the main goals of the paper, we find that human subjects are reasonably good at coping with the various sources of uncertainty. As expected, full information treatment results in the most efficient learning, and clear convergence to equilibrium predictions. Partial information (baseline) treatment leads to less efficiency, implying that players’ response to strategic uncertainty results in lower learning efficiency, and also at more conservative decisions in the SC treatment: buyers who do not know what will be the offer of their partners tend to make higher bids in order to mitigate the possibility of failing to transact. We also find that scrambled information treatment results in uniform increase of bids, in response to structural uncertainty. All in all, these findings reflect the fact that individual reaction to uncertainty results in behavior consistent with risk aversion, although the response to structural uncertainty appears to be sharper than that to strategic one, at least in our context.

In terms of learning efficiency, the efficiency of responses to uncertainty is also in line with its scale. In the BC behavior converges to Nash equilibrium, and the rate of convergence increases with the availability and accuracy of information. In NC and SC, average behavior is closer to the Nash bargaining solution. This is especially explicit in the scrambled treatment (T2), where buyers learn both less efficiently, and in a more conservative way, making increasingly larger bids in all three market types. This is consistent with risk aversion, suggesting that subjects are ready to sacrifise increasingly larger part of their expected profits in order to increase the chances of making the deal in the worst-case scenario of BC market. This behavior can also be rationalized by richer behavioral models, such as prospect theory, but the principle of Occam’s razor suggests to limit attention to the simplest scenario. With the limited feedback provided in T1, we cannot test belief learning and other strategies based on counterfactual updating. However, providing full feedback in T3 showed that subjects converged quicker towards the Nash equlibrium when they are informed of the sellers’ ask prices each trial. And even in T2, behavior was still best explained with only the outcome of the transaction. All in all, the fact that efficiency of learning increases with information implies that human bidders generally copy with structural uncertainty in a rational way.

Finally, learning strategy adopted by human subjects was primarily driven by the immediately previous feedback, and consistent with the simplest, DL rule, which was more effective than the RL algorithms. Subjects whose behavior followed more closely DL earned a higher profit at the end of the experiment. Examining the results of T2, we can conclude that this higher performance was driven more by their reaction to the trial-by-trial feedback than by game-theoretic preconceptions about optimal behavior. DL algorithms are simple: they don’t use any value function whatsoever, but simply choose between *ordered actions*. From that viewpoint, the applicability of DL is restricted to tasks where actions are structured according to an order relationship, e.g. market and auction decisions in which prices or quantities are the strategic variables. This inflexibility stands in contrast to model-free RL algorithms, which in general is always applicable, but does so at a cost of more demanding requirements imposed on information processing. Because DL algorithms parsimoniously explain bargaining decisions in which one-dimensional prices are the reference variables, whereas RL is geared to learn general values of actions or states, DL algorithms could take advantage of this informed specialization to outperform RL algorithms.

Comparison of results between T1 and T2 confirmed that although subjects factor in the degree of competition in each market type, most of their behavior can be explained by a DL algorithm, that is, regardless of the strategies specific to each market type.

## 6. Conclusions

In this study, we investigated an efficient heuristic used to optimize bargaining behavior under different types of social competition. We used a bargaining task based on a Double Auction paradigm, and an informed bargaining model which predicted behavior grounded on plausible decision-making variables which has been used to scour the brain for regions involved in learning and social aspects of bargaining behavior.

Our analysis suggests three sorts of conclusions. First, in line with our prior hypotheses, strategic uncertainty has proven to be more costly than structural one in a given market environment. At the same time, individuals are generally responsive to structural uncertainty in a rational manner: the more information about market types they possess, the more efficient is their learning, and the higher is the profit they eventually make. On the other hand, the process clearly takes time: even in T3, 24 periods were not sufficient to ensure that subjects, on average, play the Nash equilibrium. This relatively low convergence speed may be partly attributed to relatively low monetary incentives – however, even under them, individual players are able to cope with structural uncertainty efficiently.

The story of strategic uncertainty is somewhat different. In our experiment, we in most cases were unable to distinguish empirically the differences between partial (T1) and full (T3) experimental treatments, especially in comparing NC and SC markets. In the BC market, some difference exists in favour of T3 condition, which speaks in favour of the argument that subjects learn more efficiently when information about their direct competitor (another buyer) is available.

Our final result refer to learning rules. Models estimated on our data clearly speak in favour of the simplest, boundedly rational learning rules.

The DL heuristic we employ in this study is not a conventional RL rule, but a mixture of counterfactual (belief learning) and adaptive learning (Grosskopf, 2003), which incorporates knowledge of the structure of the task to make decisions. DL algorithms fared best in all bargaining tasks and markets. Despite its simplicity, implementation of DL presupposes a crucial piece of knowledge: the available bids constitutes a well-ordered set under the relation of “probability of being accepted”. DL algorithms using a simple binary learning signal rather than reward-based learning signals fit better human behavior all the experiments. Subjects whose behavior followed more closely DL also earned a higher profit at the end of the experiment. As opposed to model-free RL, DL postulates the existence of a prior knowledge about the structure of the bargaining task, namely understanding that the action values are mutually dependent through an order relationship. Finally, we showed also that constructing a utility function based on kernel density estimation of opponents behavior also didn’t fit subjects behavior satisfactorily. We reckon the reasons for this are that kernel density estimation and utility function construction are very expensive computationally, and require of a lot of data to be of practical use in uncertain and changing environments, and that their underlying assumption that opponents behave as nature precludes the consideration that mutual feedback might influence player’s behavior. On the other hand, DL makes no assumptions on the behavior of opponent players other than a minimal degree of trial-by-trial consistency and is much cheaper computationally.

## 7. Acknowledgements

We would like to thank Ellina Nurullina, Anna Drozdova, and Yulia Biryulina for assistance in recruiting subjects and collecting data, and Maria Yurevich and Karina Akopyan for providing instructions text translations.

## Appendix A. Data collection

The visual stimuli sequence were created and delivered, and presentation timings and button responses were logged by means of the software application Presentation (version 18.0, Neurobehavioral Systems, USA, www.neurobs.com). Responses were collected though three response buttons to which actions were mapped as follows: right arrow shifted cursor to right, left arrow shifted cursor to left, and spacebar confirmed bid value. Data were read in from raw files, formatted, explored, plotted and analyzed with R version 3.3.1 (R project, GNU license; R Development Core Team, 2016).

## Appendix B. Data analysis

Data were extracted, cast into data frames, reshaped, estimated, simulated, and plotted using Python and its scientific computing packages Numpy, Scipy, Matplotlib, and Pandas. All model parameters were fit employing maximum likelihood estimation. Purpose specific source code was written that created a data frame pulled from a data file, defined the functions used for estimating parameters for all the tested models, and plotted the simulated behavior of agents representing each model. Kernel density estimation was performed with a Gaussian kernel smoother (scipy.stats.gaussian_kde) with optimized kernel bandwidth. The numerical optimizer employed was a bound constrained version of the Broyden-Fletcher-Goldfarb-Shannon algorithm (L-BFGS-B). This algorithm is an implementation of a constrained optimizer of multivariate scalar functions belonging to the Python package Scipy (scipy.optimize.minimize). This optimizer was combined with a basin-hopping heuristic (scipy.optimize.basinhopping) with at least ten ‘hops’ to offset the probability that the optimizer converged into a local minimum due to the jagged geometry of the log-likelihood function.

Plotting computations for Figure 5 were performed using the plotting engine *ggplot2* (Wickham, 2008) with default parameters in the R package *stats* for the *loess* function (R Core Team, 2016).

We implemented, fitted, tested, and simulated six adaptive learning models (see Section 4) ranging from naive RL consisting in one action value for each of the possible 101 bids with two parameters, to wit, learning rate and inverse temperature; to 5-parameter DL models. The dataset consisted on the sequence of trials of all games played by the 54 subjects with the same prerecorded opponents (up to the randomization due to random-opponent matching). The same learning algorithms were fitted to the same dataset modified to comply with a coarser bid space of only 11 bins for model comparison purposes, but no substantial differences were found. Following the usual approach in this kind of estimation problems (Daw et al., 2006), first parameters were yoked (fixed) across all market types and subjects and optimized in a global objective likelihood function encompassing all subjects (Table 4, FFX column):

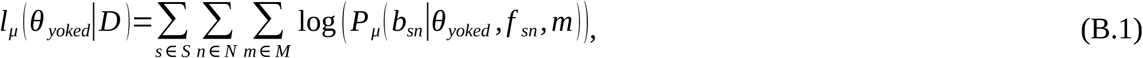

where l_μ_ is the log-likelihood function for model μ, θ_yoked_ is the parameters vector of model μ (for example, for naive RL θ_yoked_=(α,β)), P)), P_μ_ is the likelihood of model μ choosing a specific bid given parameters θ and feedback f_sn_, *s* is the index ranging over the subjects, *n* is the index of the block, and *m* is the index for market competition type.

This reduces parameter estimator variances at the cost of losing the ability to make between-subject parameters comparisons by conflating between-subject with within-subject variance. Given the scarcity of within-subject samples and the jagged geometry of the resulting objective functions, it has the advantage of having less variance at the expense of some bias. The alternative of running individually the numerical optimizer for each subject leads to poor optimization convergence and trapping in suboptimal local minima for some subjects (see Table 4 legend). Nevertheless, in order to confirm that the fitted parameters were reasonable, an additional fitting routine was accomplished individually for each subject, and between-subject standard errors of the mean parameter values and BIC scores were calculated too for each subject *s* (Table 4, RFX column):

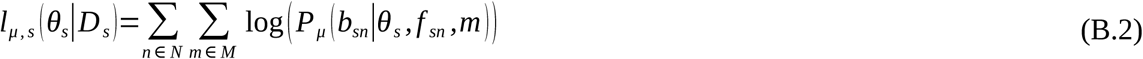

Code implementing artificial bidders, model fits, and simulation results is available under the MIT license on the hosting service GitHub (https://github.com/mmartinezsaito/adaptive-learning-in-auctions).

## Appendix C. Theoretical backgroupd

### Equilibrium predictions

Because all valuations in our experiment are common knowledge, players in each market type face a normal form game of complete information. In NC, single buyers and sellers play a coordination game. Price *p* for NC is determined as follows: if *s*_1_≤*b*_1_ then *p*=*b*_1_, while if *s*_1_>*b*_1_, then no trade occurs (the seller asks more than the buyer bids). Payoffs are 10-*b*_1_ and *b*_1_ to buyer and seller, respectively, if *b*_1_ ≥*s*_1_, and 0 otherwise. Hence, the set of equilibria consists of all pairs of strategies *s*_1_=*b*_1_, s.t. *s*_1_+*b*_1_=10. See Table 1 for a simplified payoff matrix^1^.

**Table 1.**
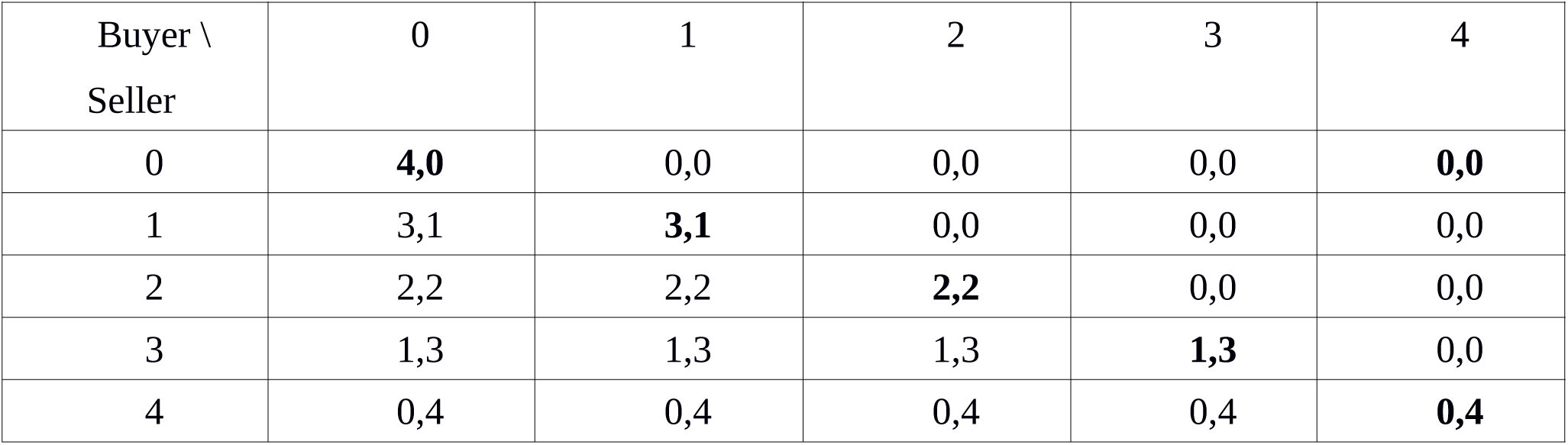
Simplified payoff matrix and equilibria in the NC market type with 5 bids / ask prices (0, 1, 2, 3, 4) instead of the 10 of the actual task. Payoffs in Nash Equilibrium profiles are highlighted in bold.

| Buyer \ Seller | 0 | 1 | 2 | 3 | 4 |
| --- | --- | --- | --- | --- | --- |
| 0 | **4,0** | 0,0 | 0,0 | 0,0 | **0,0** |
| 1 | 3,1 | **3,1** | 0,0 | 0,0 | 0,0 |
| 2 | 2,2 | 2,2 | **2,2** | 0,0 | 0,0 |
| 3 | 1,3 | 1,3 | 1,3 | **1,3** | 0,0 |
| 4 | 0,4 | 0,4 | 0,4 | 0,4 | **0,4** |

<sup>1</sup> . This game also has an isolated corner equilibrium with minimum bid for the buyer and maximum for the seller. Unlike the others, this equilibrium is not perfect, so in the future we do not consider it.

In BC, transaction takes place with the buyer who made the largest bid, so *p*=max(*b*_1_,*b*_2_) if *p*>*s*_1_ (in particular, it is possible that *p*=*b*_1_>*s*_1_>*b*_2_). Payoff to the higher-bidding buyer is *10-p*, to the seller *p*, and to the lowest-bidding buyer *0*. The buyer who made the strictly smallest bid gained nothing. The basic prediction for BC is that of Bertrand competition: buyers shall drive the price up to the maximum, with expected payoff of 0, while the single seller will gain *10*.

Finally, in SC, price *p*=*b*_1_, i.e. equals buyer’s bid, provided *b*_1_>min(*s*_1_,*s*_2_). If only one seller offers a good for sale at a price *s*_1_<*b*_1_, while the other asks *s*_2_>*b*_1_, transaction takes place with the former seller; if both *s*_1_ and *s*_2_<*b*_1_, the seller who transact with the buyer is chosen at random (reasons for that setup shall be explained shortly). No seller has incentives to increase its bid, while decreasing it results in Bertrand competition with equilibrium *p*=0: payoffs to both sellers is *0*, while that of the buyer, *10-p=10*.

#### C.2 Treatments effects

*Proposition 1*. A risk averse individual should increase bids in the T2 treatment (T2) in comparison to T1 and T3 treatments.

Proof: Denote the probability of market types B,C,S in T2 as *q*_*B*_, *q*_*N*_, *q*_*S*_, resp. Utility of getting a transaction from bid *b* in equilibrium is

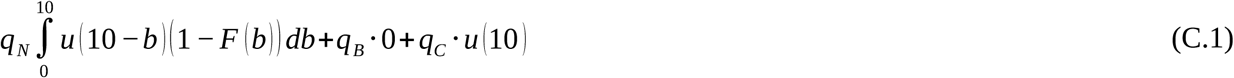

In case of risk neutrality, this expression amounts to

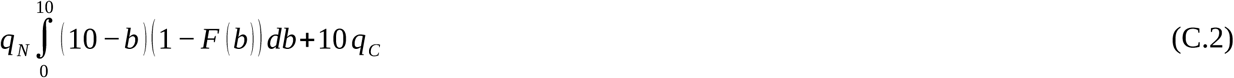

Since a linear combination of two concave functions is concave, the certainty equivalent of (1) is less than that of (2). Accordingly, the maximand of (1), or optimal value of b under risk neutrality should be higher than that under (2).

*Proposition 2*. Risk averse individual should make larger bid in SC than risk neutral invidudal.

Proof: The expected utility of getting the transaction at any given bid b is given by

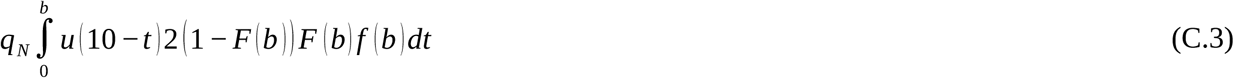

where the probability density function comes from the second-order statistic (Krishna, 2010, p.282). Any risk-averse individual has convex utility function, resulting in lower certainty equivalent, and hence should make lower bid than the risk-neutral counterpart.

Krishna, V. (2010) Auction Theory. 2^nd^ edition, Academic Press.

## Appendix D. Task instructions translated to English

Dear participant,

During this experiment you will be asked to make decisions in a economic game. Each decision will lead to a particular financial result.

You are the owner of a fish restaurant, who goes to the market to buy some fish. Each trial you are given 10 monetary units on credit without charge for interest. You may spend any amount of your money on fish. You always buy the same amount of fish (in kilograms). Then you sell the fish in your restaurant for 10 monetary units. The less money you spend on fish, the more income you get by selling it in the restaurant.

You get your income as a cash bonus (see below). If you refuse to buy fish in a single trial, the borrowed money will be automatically returned without interest. This means that you will neither earn nor lose money in this trial. You will face three typical market situations.

### Market situations

Market situations will vary depending on the quantity of sellers and buyers on the market. First of all, there may be either one or two sellers on the market. Secondly, you may be the only buyer on the market or there may be another one. During each trial (one financial operation) you will be informed about the quantity of buyers and sellers. The sellers and buyers have participated in previous sessions of the experiment (see section “players”). Each trial players are selected randomly from the database consisting of 16 sellers and 16 buyers (participants in previous experimental sessions).

You will not recognize the participants you will be playing with.

At the end of the game the participants who played with you will get extra cash bonus which depends on the results of the financial operations during the game (see more in section “payment”).

Remember, that you are going to play with real participants and that the experiment meets the standards of the HSE ethics committee.

Three possible market situations:

There is only one seller on the market and you are the only buyer.

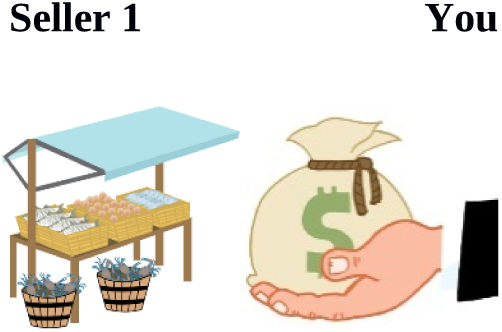
There are two sellers on the market and you are the only buyer.

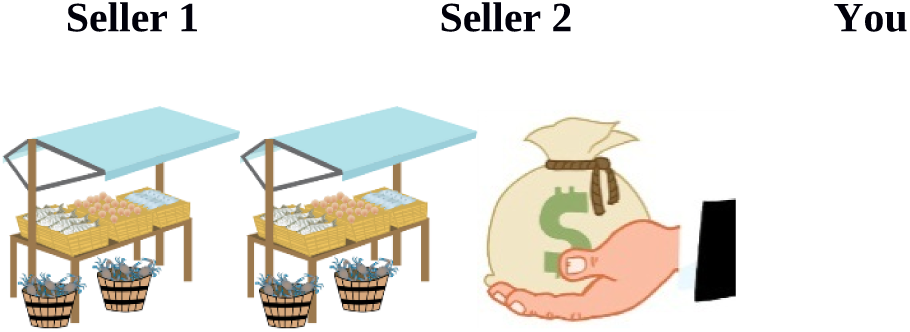
There is only one seller on the market and another buyer.

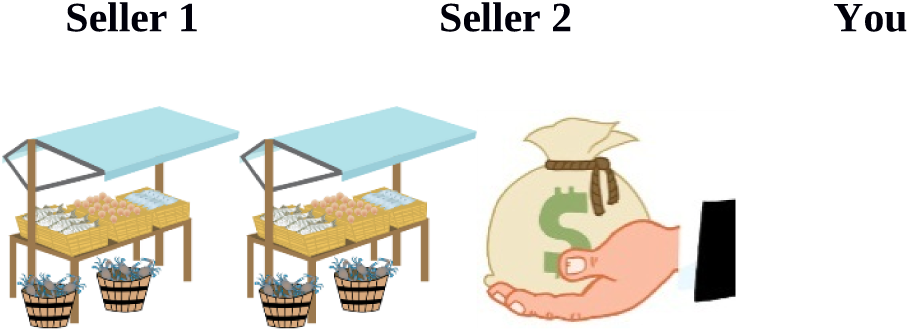
Keep in mind that the transaction may occur only between two people (the buyer and the seller).
If both sellers accept your offer, one seller is selected randomly and the other one doesn’t sell fish and doesn’t make a profit.
If both you and the other buyer make acceptable offers, the seller makes a deal with the one who offers a higher price. In other words, the buyer who suggested lower price doesn’t buy fish in this trial and doesn’t make a profit. If both buyers make the same offer, one of them is selected randomly.

### Each trial consists of three stages: firstly, you are informed about the market situation. At the second stage you make a financial proposal to a seller or two sellers. At the third stage you find out the results of the deal

***Stage 1*** *Market situation*

***Stage 2*** *Proposal*

After you are aware of the market situation, you are about to make a proposal to one or two sellers. You may spend any amount of money from 0 M.U. to 10 (as you only have 10 M.U.), this means that your proposal may also be: 0, 0.1, 0.2, 0.3, … 9.7, 9.8, 9.9, 10 M.U.

Each trial sellers establish the floor price for fish while you make your decision. No one, except for the sellers, knows the floor price. The floor price is the lowest acceptable one, so if you propose a higher or equal price, one of the sellers will accept the proposal. Thus, you will buy fish at your price. If neither seller accepts your proposal you are not going to buy fish and make a profit during the trial.

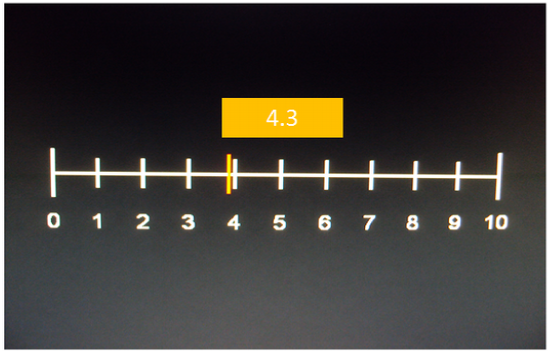

***Stage 3*** *Acceptance of proposal*

If a seller accepts your proposal and you buy the fish, you are to sell it in your restaurant for 10 monetary units. Still you have to give back the 10 M.U. that you have lent. Remember, that the money left after the fish purchase is your profit during the trial (10 – fish price = profit). The seller’s profit is the money you’ve paid for the fish.

*Examples:*

You’ve lent 10 monetary units and bought the fish for 4.3 M.U. and so you have 5.7 M.U. left (10 – 4.3 = 5.7). You sell the fish in your restaurant for 10 M.U. and repay 10 M.U. This means that the money left after the fish purchase is your profit (5.7 M.U. in this example).

If you buy fish for 8 M.U. your profit is 2 M.U. (10 - 8 = 2).

If you buy fish for 2.5 M.U. your profit is 7.5 M.U. (10 – 2.5 = 7.5).

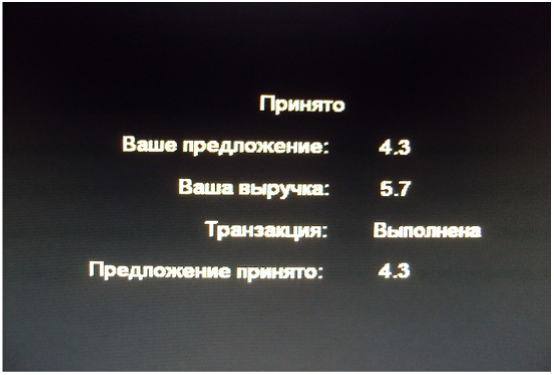

*Rejection of proposal:*

If your proposal is lower than the floor price and neither seller accepts it, you just give 10 M.U. back and don’t make a profit during the trial. If another buyer proposes a higher price, the seller will accept it and reject your proposal. In this case you will not get profit. Sellers don’t make a loss during the experiment, so if a seller rejects both offers they will not get a profit.

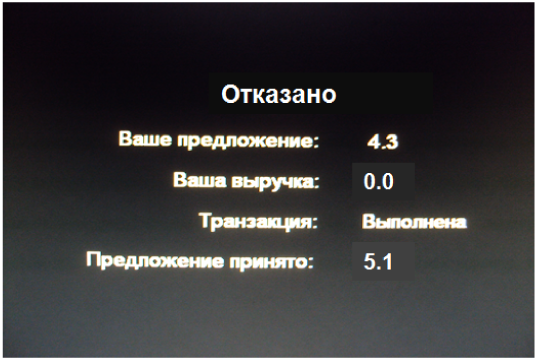

### Experiment demo

The pictures below illustrate 3 types of slides you will see during the experiment.

***Stage 1: market situation***

**Seller**

You will see if there is only one or there are two sellers on the market. If there is a “+” sign, then this seller is present; in case of “–” sign, the seller is absent. The situation with two sellers and one buyer (you) is illustrated in the picture.

**Buyer**

If there is “+” sign, then this buyer is present. According to this picture you are the only buyer.

***Stage 2: proposal***

Here you can establish your price for the fish. Press button 1 to move the cursor (red line) to the left and button 2 to move it to the right. The chosen value is shown in a bar above the scale (in this example the value is 4.3). To confirm the choice, please, press button 3. You are to make a decision in less than 5 seconds.

***Stage 3: feedback***

**Acceptance/rejection**

Shows whether your proposal was accepted or not.

**Your proposal**

Shows your proposal to the seller.

**Your income**

Shows your profit on the operation. In this case it is – 5.7 M.U. as the proposed and accepted price was 4.3 M.U. If your proposal is rejected, the income is 0.

**Transaction**

Displays whether the transaction was committed or not. If your proposal was accepted, you will see that the transaction was committed. <u>This information is especially important if there is another buyer on the market and your proposal was rejected. In this case the sign informs that the transaction was committed by the other buyer.</u>

**Accepted proposal**

Shows the price accepted by the seller. This information might be helpful in case there is another buyer and your proposal was rejected.

**Please, notice** that in this example your proposal was rejected and the seller accepted another proposal of 5.1 M.U. because it was larger than yours.

### Payment for participation in the research

You will be payed 50 rubles for participation and an extra bonus depending on your decisions. The bonus depends on the results of your deals during the experiment. One trial among each market situation will be selected (this may be the case when you didn’t go to the market and received 0 income). The income of three selected trials and the bonus are summarized: 1 M.U. equals 5 rubles. Thus, you can get from 0 to 150 rubles in cash bonus which in sum with the initial payment (50 rur) gives from 50 to 200 rur.

### Duration of the experiment

There will be 72 trials in the experiment which will take you about 30 minutes.

If you have any questions, please, ask the researcher now.

You will have a training session in the beginning of the experiment.

Thank you for your participation!

## Footnotes

1 This game also has an isolated corner equilibrium with minimum bid for the buyer and maximum for the seller. Unlike the others, this equilibrium is not perfect, so in the future we do not consider it.

